# Comparative Extraction of Cellular Features from High-Resolution Volume Imaging

**DOI:** 10.1101/2025.02.01.636042

**Authors:** Yuko Mimori-Kiyosue, Tomoko Hamaji, Akiko Hayashi, Miwa Okada, Koichiro Higasa, Kazuyuki Kiyosue, Yuki Gomibuchi, Takuo Yasunaga, Takashi Washio, Satoshi Hara

## Abstract

High-resolution volume imaging techniques, such as lattice light-sheet microscopy (LLSM), generate vast and complex datasets that demand advanced analytical approaches to uncover biologically meaningful insights. While LLSM’s high spatial and temporal resolution provides critical data for understanding cellular processes, distinguishing subtle differences between cells in distinct states remains challenging. Here, using an adaptive, human-interpretable kernel-based calculation strategy, we developed Machine-Learning-Based Visual Extraction of Structural Features (M-VEST), a method designed to identify and interpret structural differences with high precision. By applying M-VEST to mitotic cells, we uncovered novel functions of the oncogene Aurora kinase A, demonstrating its utility in revealing previously undetected features. Validated using LLSM datasets, M-VEST offers a scalable framework for analyzing large and complex imaging data, advancing insights into cellular dynamics and beyond.

## Introduction

Recent advances in imaging technologies, particularly light-sheet microscopy, have revolutionized 3D volume imaging by capturing detailed datasets of biological processes in unprecedented resolution (1). Among these, lattice light-sheet microscopy (LLSM) achieves diffraction-limited spatial resolution but generates vast and complex datasets that are challenging to analyze manually (2, 3). While machine learning (ML) has been effectively applied to technical tasks such as image enhancement, super-resolution, and segmentation (4–8), extracting biologically meaningful insights, particularly through comparative analysis, remains a significant challenge.

This difficulty is evident in the case of LLSM, where each experiment can produce datasets ranging from several hundred gigabytes to terabytes, creating considerable challenges in both storage and computational processing (9). The obstacles stem from the immense data volume, requiring high-performance computing resources for efficient processing, and the limitations of current ML models, which struggle to adapt to the complexity and variability inherent in biological structures. Moreover, the lack of standardized workflows for analyzing high-resolution 3D datasets further complicates the extraction of meaningful insights. Comparative analysis, in particular, involves identifying and interpreting subtle, context-dependent variations in multidimensional data, such as differences between cellular states or experimental conditions.

To address these challenges, we developed an adaptive and interactive ML-based method named <u>M</u>achine-Learning-Based <u>V</u>isual <u>E</u>xtraction of <u>S</u>tructural Features (M-VEST). This method identifies structural differences by selecting optimal kernel sizes and corresponding weights tailored to diverse biological structures, enabling precise feature extraction. M-VEST is particularly effective in adapting to complex datasets and detecting subtle structural differences. When applied to two critical components during cell division—microtubules and chromosomes—it revealed new insights into the role of the oncogene Aurora kinase A (AURKA), a key regulator of mitosis frequently overexpressed in cancer (10). These findings underscore the potential of M-VEST to uncover novel biological insights through advanced imaging data analysis.

## Results

### M-VEST Workflow: Optimizing Feature Extraction for High-Resolution Volume Imaging

The M-VEST workflow consists of four main phases: 1) model training, where experimental image data from two distinct groups (e.g., images from different phases of cell division) are used to optimize the model; 2) applying the trained model to a test dataset; 3) model validation and tuning of the learning process; and 4) analyzing the generated feature maps (Fig. 1 and Fig. S1).

**Fig. 1.**
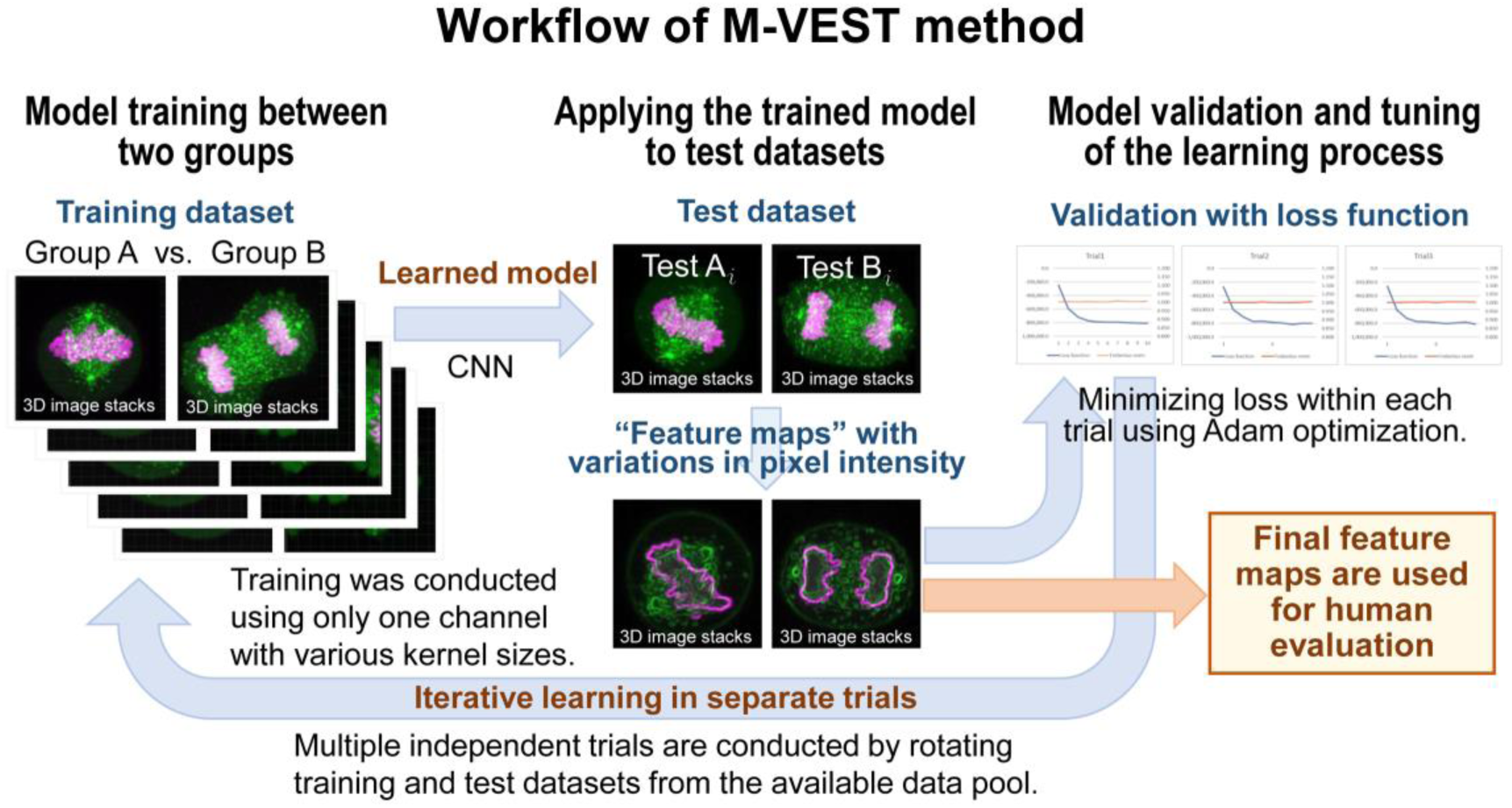
Workflow of the M-VEST Method. Feature maps generated after training highlight pixel intensity variations, which are used to interpret biological results. See also Fig. S1 and Methods. Note: the data comprise two-channel multistack and multipoint images, but training is performed on each slice of a single channel.

Initially, 3D image stacks from two distinct groups are processed through convolutional neural networks (CNNs), where various cubic kernel sizes are used to determine the optimal size for capturing features that differentiate the groups. During training, the convolutional filter weights are optimized for each kernel size to enhance the detection of specific structural features. Following training, the model is applied to the test datasets, and feature maps are generated to evaluate the model’s performance through a loss function. These feature maps, produced by applying optimized convolutional filters, highlight relevant structural features in the data. Model optimization is achieved using the adaptive moment estimation (Adam) algorithm (11) and a combined loss function that integrates the standard entropy loss with a custom regularization term based on the Frobenius norm (12). Once the model demonstrates satisfactory performance on the test dataset, the generated feature maps can be examined by human experts. These maps provide clear, human-interpretable representations of extracted features, facilitating intuitive analysis of structural differences within the observed data.

In this study, we used timelapse images of mitotic HeLa cells collected with LLSM (XYZ resolution: 320×320×∼370 nm for GFP) to analyze the behavior of microtubules and chromosomes (3); however, the data were not treated as a time series. HeLa cells expressing the microtubule plus-end marker EB1-GFP and the chromosome marker H2B-RFP (A1 cells) (3) and A1 cells overexpressing AURKA by approximately four times the endogenous level (AURKA cells) (13) were used to compare different cellular states. Prior to the main M-VEST analysis, we optimized the image preprocessing by creating one dataset with brightness normalization (“normalized”) and another with background brightness removal (“mask + background threshold”) (Fig. S2, Table 1). While the normalized data showed suboptimal learning performance, the mask + background threshold dataset (cubic kernel size 5×5×5, i.e., 0.5 µm because 1 pixel = 0.1 µm) resulted in optimal performance for EB1-GFP, while a kernel size of 7×7×7 was optimal for H2B-RFP (Fig. S3; see also Supplementary Results and Discussion).

### Feature Map Analysis of EB1-GFP Intensity in Metaphase and Anaphase Cells

We first compared the EB1-GFP channel in A1 metaphase and anaphase (Fig. 2a, top). Metaphase is marked by chromosome alignment at the equatorial plane, while anaphase begins with chromosome separation. EB1-GFP binds to the plus end of growing microtubules, forming a “comet-like” distribution (14) that reflects the GTP cap, a region of GTP-bound tubulin, serving as a proxy for GTP cap size (15). The GTP cap length depends on microtubule growth speed, with faster growth yielding longer caps (15–17). In optical microscopy, this distribution appears as structures 0.3–1 µm in length due to fluorescence spread and resolution limits (14, 18).

**Fig. 2.**
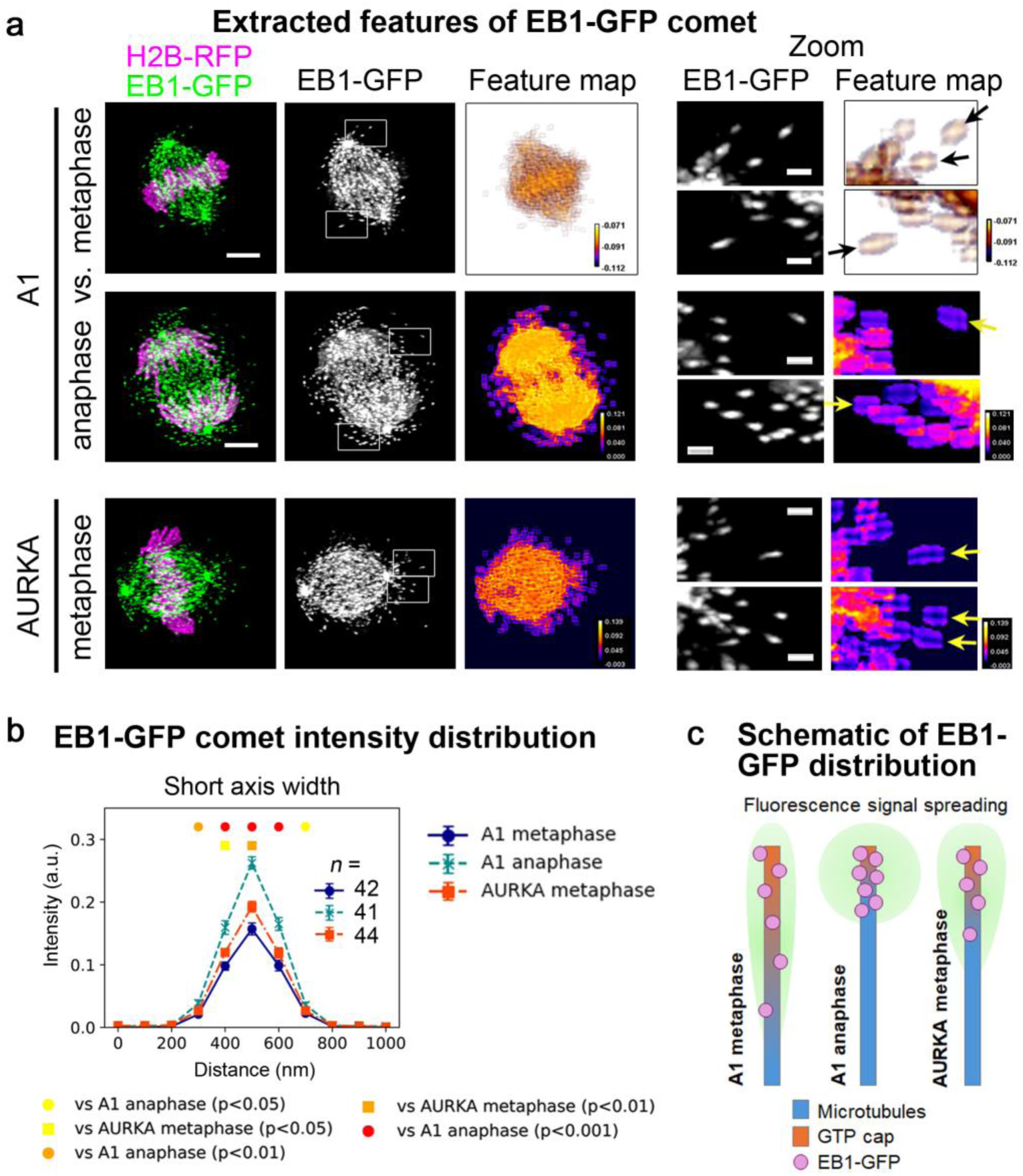
Analysis of EB1-GFP comet image features and intensity distribution. (**a**) Extracted features of EB1-GFP comets. Representative images of HeLa cells expressing EB1-GFP and H2B-RFP with/without AURKA (A1 and AURKA cells) and their corresponding feature maps. Only AURKA metaphase is shown below because A1 metaphase results are similar in both comparisons. The boxed areas are zoomed for detailed visualization. See also Movies S1–S4. Black and yellow arrows indicate examples where comets appear brighter along the long axis in A1 metaphase and darker in A1 anaphase and AURKA metaphase, respectively. Scale bars: 5 µm. (**b**) Graphs showing the intensity profiles of EB1-GFP comets along the short axis width. Each plot compares A1 metaphase (blue), A1 anaphase (green), and AURKA metaphase (orange) cells, with intensity plotted against distance (nm). Statistical significance was assessed using Welch’s t-test (unequal variance t-test) at each position, comparing A1 metaphase with A1 anaphase and AURKA metaphase. Significant differences are marked by colored dots along the x-axis: yellow for *p* ≤ 0.05, orange for *p* ≤ 0.01, and red for *p* ≤ 0.001. Error bars represent the standard error (SE) at each data point. (**c**) Schematic illustration of EB1-GFP distribution under different conditions. The diagram visualizes fluorescence signal spreading and its relationship to microtubules (green), GTP caps (orange), and EB1-GFP (purple) in each condition, highlighting differences in comet size and signal distribution.

Movie 1 shows 3D stack images, while Movie 2 shows 2D projections over time. Both movies include the test images used for model evaluation on the left and their corresponding feature maps generated by the ML model. These visualizations illustrate how the model processes the data to highlight structural differences relevant to cellular states. In feature maps highlighting phase-specific pixels, metaphase comets have a brighter, elongated axis (Fig. 2a, arrows), indicating that longer comet structures are characteristic of metaphase. This observation aligns with the reported faster average microtubule growth speed, as the microtubule growth rate averaged 0.449 µm/s in metaphase and 0.346 µm/s in anaphase (Fig. S4a, b) (2).

While metaphase comets appear brighter and are elongated along their axis, the sides of the comets are darker in metaphase and brighter in anaphase—an unexpected observation. This difference in lateral intensity suggests that the EB1-GFP signal is more broadly distributed in anaphase, implying a higher molecular density. Although the knowledge that differences in microtubule growth speed alter EB1 density has not been reported before, careful measurements of our A1 metaphase and A1 anaphase data confirmed that the EB1-GFP intensity distribution in anaphase was, on average, 1.6 times wider and brighter than in metaphase (Fig. 2b, Fig. S4c). The mechanisms underlying this observation are discussed in detail in the Supplementary Results and Discussion. This result demonstrates M-VEST’s ability to capture subtle structural differences that may have previously gone unnoticed. Furthermore, it highlights the utility of analyzing the lateral intensity distribution of EB1-GFP comets as an effective indicator of microtubule growth-dependent changes in EB1-GFP density.

### Comparative Intensity Analysis of EB1-GFP Images in Control and AURKA-Overexpressing Metaphase Cells

Next, we compared EB1-GFP in A1 and AURKA-expressing metaphase cells (Fig. 2a, bottom). AURKA’s activation at centrosomes contributes to the maturation of these structures, increasing their ability to nucleate microtubules (19). The M-VEST analysis revealed that AURKA-expressing cells exhibit brighter comet sides, suggesting slower microtubule growth, which can be interpreted as brighter sides correlating with slower growth rates, as observed as a characteristic of A1 anaphase (Fig. 2b, Fig. S4c). AURKA-expressing cells had 1.4 times the number of EB1-GFP comets observed in A1 metaphase cells (average of 807/cell versus 583/cell) (13), suggesting a corresponding increase in the number of microtubules in AURKA-expressing cells. Given that an increased number of microtubules depletes cytoplasmic tubulin, thereby slowing the growth rate (20), the interpretation from the M-VEST analysis aligns with these known facts, further validating its ability to capture key features of microtubule dynamics.

Measurements of the EB1-GFP comet signal width in the images used for the ML analysis indicated that, on average, the signal width in A1 anaphase is approximately 151 nm greater than in A1 metaphase, and the signal width in AURKA metaphase is approximately 80 nm greater than in A1 metaphase (Fig. S4c). The voxel dimensions are 0.1 µm on each side in the images, so the ML model is sufficiently sensitive to capture subpixel variations. However, individual EB1-GFP comet images are likely to exhibit greater width variations than the average value (Fig. S4a, b). These individual variations may contribute to the ML model’s ability to detect subtle structural differences at almost single-pixel level as it learns to generalize from a range of comet widths. In this way, the ML model may capture a broader structural profile by accounting for individual comet characteristics, thus enhancing its sensitivity and accuracy in differentiating cellular states. These results are consistent with the ML-based detection of mitotic cells in pathological specimens, where incorporating subtle diversity within the data improves sensitivity and accuracy (21).

### Structural Changes in Chromosomes Driven by AURKA Overexpression

Finally, we analyzed the chromosome (H2B-RFP) channel in A1 and AURKA metaphase cells, focusing on the structural changes in thread-like chromosomes (approximately 1 µm thick) in relation to AURKA overexpression. The feature map predicted the presence of an additional structure outside the AURKA chromosome, suggesting that the chromosome is thicker in AURKA cells (Fig. 3a). No prior evidence directly links AURKA to the regulation of chromosome thickness, but studies suggest that histone acetylation can influence this characteristic (22). Fluorescent immunostaining of major acetylated histones—H2B, H3, and H4—revealed a significant increase in H2B acetylation alone (Fig. 3b). This is particularly relevant because the phenomenon of histone-type-specific acetylation enhancement has only recently been understood. H2B-specific acetylation enhancement can indeed occur (23), suggesting that AURKA may act in concert with the CBP/p300 pathway to selectively enhance H2B acetylation. Given that alterations in chromosome thickness can affect transcriptional activity, we hypothesize that AURKA overexpression in cancer might influence transcription. Furthermore, microtubule– chromosome interactions are apparently affected (Fig. 2c), further supporting our interpretation of the feature map, because previous observations indicate that when chromosomes expand, microtubules may fail to pause on the chromosomes and instead pass through them (22).

**Fig. 3.**
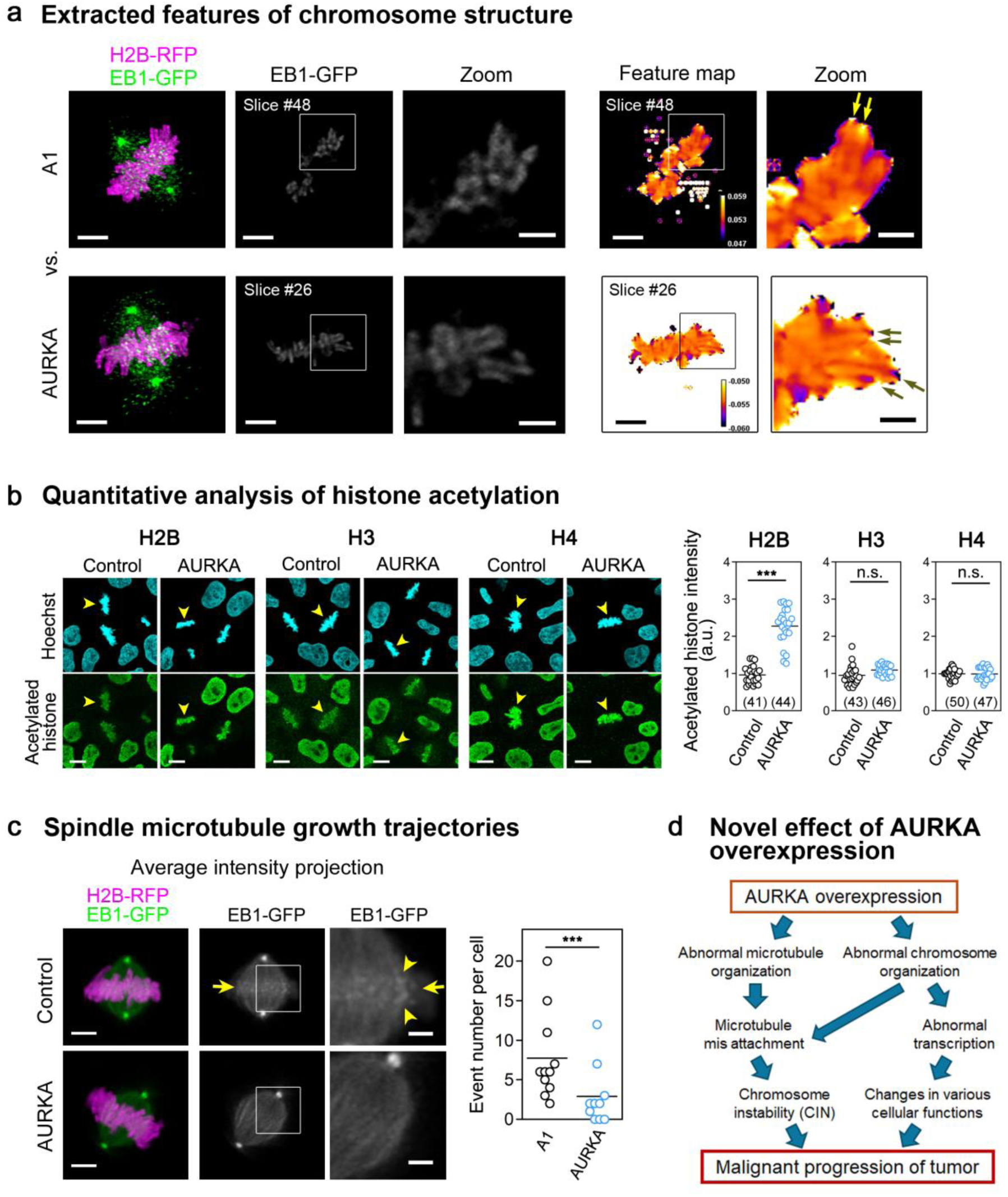
Effect of AURKA overexpression on chromosome structure, histone acetylation, and spindle microtubule dynamics. (**a**) Features of chromosome structure extracted using M-VEST. Representative images showing EB1-GFP and H2B-RFP in A1 and AURKA cells are presented as slice images. Feature maps and zoomed views highlight differences in chromosome structure between A1 and AURKA metaphase cells. See also Movies S5 and S6. Scale bars: 5 µm. (**b**) Histone acetylation levels in control and AURKA-expressing cells. Immunofluorescence of acetylated histones (H2B, H3, and H4) with Hoechst staining highlights chromosomes in mitotic cells (arrowheads). Quantitative analysis reveals significant differences in H2B acetylation levels between control and AURKA cells (t-test, \*\*\**p* < 0.001), with no significant changes observed for H3 and H4 (n.s. = not significant). Sample sizes are indicated in parentheses. Scale bars: 5 µm. (**c**) Spindle microtubule dynamics visualized through EB1-GFP intensity projections. Average spindle microtubule growth trajectories in A1 and AURKA-expressing cells are shown. Arrows indicate the equatorial plane where kinetochores align on the spindle, and arrowheads mark potential microtubule–chromosome attachment sites (25). Quantification of dot number per cell shows a significant decrease in AURKA-expressing cells compared to controls (t-test, \*\*\**p* < 0.001). Scale bars: 5 µm. (d) Summary of AURKA-induced cellular changes. A schematic illustrates the novel effects of AURKA overexpression identified in this study, including alterations in chromosome structure, histone acetylation, and spindle microtubule dynamics. Sample sizes for A1 cells and AURKA cells are 11 and 10, respectively.

The findings from M-VEST highlight previously uncharacterized roles of AURKA, including its ability to slow microtubule growth, alter microtubule–chromosome interactions, and potentially regulate transcription by modulating chromosome thickness (Fig. 3d). These results underscore the potential of M-VEST to identify subtle structural changes in cellular processes that may otherwise remain undetected. Further exploration of the generated feature maps could uncover additional biological insights, paving the way for deeper understanding of chromosomal dynamics and their implications in cancer.

## Discussion

The findings presented here highlight the effectiveness of the M-VEST method in elucidating molecular functions by detecting subtle structural variations with single-pixel-level sensitivity. This capability is largely attributable to the high 3D resolution of LLSM, which, when combined with M-VEST, facilitates an adaptive approach to studying cell dynamics. By enabling precise feature extraction, M-VEST effectively captures complex biological structures and processes, offering new perspectives on cellular architecture and function.

A key advancement demonstrated by M-VEST lies in its application to mitotic processes, particularly in analyzing the role of AURKA. For instance, M-VEST revealed changes in EB1-GFP comet width associated with alterations in microtubule growth rate driven by AURKA overexpression. This finding highlights M-VEST’s ability to detect subtle structural variations and offers new insights into the dynamic regulation of microtubules during mitosis. Furthermore, M-VEST uncovered variations in chromosome thickness associated with AURKA overexpression, suggesting new dimensions in chromosomal dynamics and histone modifications that could influence transcriptional regulation. These findings position M-VEST as a pivotal tool in modern biological imaging, bridging the gap between advanced microscopy and ML.

M-VEST’s kernel-based approach contributes significantly to its scalability and adaptability. Unlike traditional methods that rely on fixed parameters, M-VEST dynamically selects optimal kernel sizes tailored to diverse biological structures, ensuring precise feature extraction across varying scales. For instance, M-VEST successfully differentiated EB1-GFP comets in metaphase and anaphase, highlighting subtle variations in comet width that are often overlooked by other methods. Moreover, its adaptability enables application across multiple imaging platforms, including super-resolution microscopy and electron microscopy, broadening its potential for investigating distinct biological systems. This kernel-based flexibility also facilitates handling large datasets, as the computational efficiency of a single-layer CNN minimizes resource demands while maintaining high sensitivity. As such, M-VEST is uniquely positioned to address challenges posed by complex and high-dimensional imaging data, making it an indispensable tool in modern biological imaging.

Despite these advancements, interpreting the biological significance of M-VEST-generated feature maps requires specialized knowledge, particularly in the context of subtle structural differences. This underscores the importance of refining workflows to enhance interpretability and expand accessibility. Currently, M-VEST relies on a single-layer CNN for feature extraction, which has proven effective for the current scope of analysis. However, incorporating deeper CNN architectures or advanced methodologies, such as transformers (24), could enhance the method’s ability to capture intricate and hierarchical structural details. These developments, while promising, would necessitate additional computational resources to manage the increased data complexity and processing requirements. Additionally, beyond high-precision systems like LLSM, M-VEST has the potential to be adapted to various imaging platforms and data types. This adaptability could uncover biological mechanisms that are not readily observable with high-resolution imaging alone, broadening the horizons of cellular and molecular biology research.

Overall, M-VEST represents a transformative approach to integrating ML with advanced imaging technologies, enabling the investigation of complex cellular dynamics. Its demonstrated ability to uncover subtle structural features not observable with conventional methods highlights its potential to accelerate discoveries in cell biology. As M-VEST continues to evolve through methodological refinements and expanded applications, it promises to unlock new dimensions of biological understanding in both research and clinical contexts.

## Supporting information

Supplementary Results and Discussion

## Acknowledgments

We thank Drs. Wesley Legant, Bi-Chang Chen and Eric Betzig (Janelia Research Campus, HHMI) for generously providing LLSM data for this study. We are also grateful to Dr. Marija Zanic (Vanderbilt University) for her insightful guidance on microtubule dynamics analysis, to Dr. Gokul Upadhyayula (University of California, Berkeley) for his valuable comments and advice on image processing, and to Dr. Tomoya Kitajima for his helpful advice on chromosome analysis. The computational resources of the AI Bridging Cloud Infrastructure (ABCI) provided by the National Institute of Advanced Industrial Science and Technology (AIST) were used.

## Funding

Y.M.-K. was supported by the Japan Society for the Promotion of Science– NEXT program (LS128), the Takeda Science Foundation, the Uehara Memorial Foundation, a Grant-in-Aid for Challenging Research (Pioneering) (JSPS KAKENHI Grant Number 20K20379), Japan Science and Technology Agency Core Research for Evolutional Science and Technology (no. JPMJCR1863), and an intramural grant from RIKEN. T.W. was supported by Japan Science and Technology Agency Core Research for Evolutional Science and Technology (No. JPMJCR1666).

## Author contributions

Y.M.-K., T.H., A.H. and M.O. performed experiments. K.H. carried out computational calculations. Y.M.-K., K.K., Y.G. and T.Y. performed image data analysis and statistical data analysis. T.W. and S.H. gave conceptual and technical advice. Y.M.-K. conceived and managed the project, performed experiments, and wrote the manuscript.

## Competing interests

The authors have no competing interests to disclose.

## Notes

### Competing Interest Statement

The authors have declared no competing interest.

