## Supplementary Results and Discussion for "Comparative Extraction of Cellular Features from High-Resolution Volume Imaging"

Yuko Mimori-Kiyosue *et al.*

**This PDF file includes:**

Supplementary Results and Discussion

Supplementary Materials and Methods

Supplementary References

Figs. S1 to S4

Legend for Tables S1 and S2, and Movies S1 to S6,

**Other Supplementary Materials for this manuscript include the following:**

Tables S1 and S2

Movies S1 to S6

### Supplementary Results and Discussion

#### Optimization of preprocessing and kernel sizes in machine learning model training

In this study, we focused on two key structures during cell division: the growing ends of microtubules and chromosomes. These two structures present different challenges in image analysis. The EB1-GFP signals represent small, scattered comet-like structures, typically ranging from 0.3–1 μm long in fluorescent microscpy, dispersed throughout the cell. In contrast, the H2B-RFP channel captures thread-like chromosomes, approximately 1 μm thick, that are grouped and concentrated in a specific region of the cell. Optimization of preprocessing methods was crucial for extracting information from biological images with poor signal-to-noise ratios and high noise levels, as well as issues related to brightness fluctuations, fading, and variations in brightness across different cells. In addition, kernel size optimization was vital in ensuring accurate machine learning (ML) model training, as both preprocessing and kernel size significantly influence the outcome.

Two preprocessing methods were applied: brightness normalization and high-thresholding (Fig. S2a). During the evaluation of ML conditions, when using images processed with normalization only, the learning curve did not converge sufficiently well and the reproducibility was poor (Fig. S2b, normalized). Because the brightness in the background of the target structure could be the cause, we decided to perform advanced processing to remove the background brightness. Specifically, after normalization, a mask was used to set the extracellular regions to zero, followed by high-thresholding to remove diffuse cytoplasmic signals (mask + background threshold).

Figure S2a shows representative preprocessed images of A1 metaphase, A1 anaphase, and AURKA metaphase cells, illustrating the two preprocessing methods. The top row shows images for both EB1-GFP (green) and H2B-RFP (magenta) channels, with both preprocessing methods applied and displayed across the brightness range of 0–1. The middle and bottom rows present the EB1-GFP and H2B-RFP channel images, respectively, each showing the patterns of background brightness across different display ranges to help visualize how the background signal was treated for each condition.

Figure S2b compares the learning curves during 3D model training across different kernel sizes in both the EB1-GFP (green) and H2B-RFP (magenta) channels. For simplicity, the kernel sizes are referred to as 3, 5, 7, 9, and 11 to represent their full 3×3×3, 5×5×5, …. form. For the EB1-GFP channel, comparisons are made between A1 metaphase and A1 anaphase. In both conditions, kernel sizes of 3, 5, 7, 9, and 11 are compared across trials in the left plot, followed by kernel size 5 as a representative example in the adjacent plot. For the A1 metaphase vs AURKA metaphase comparison, based on the results of the A1 metaphase vs A1 anaphase analysis, only kernel sizes 3, 5, and 7 were tested, and the loss values reflect consistent image quality across these conditions. For the H2B-RFP channel, comparisons are made between A1 metaphase and AURKA metaphase. In both the normalized and mask + background threshold conditions, kernel sizes 5, 7, 9, and 11 are compared across trials in the first plot, followed by kernel size 7 as a representative example in the adjacent plot.

The learning curves illustrate the relationship between the number of epochs and the corresponding loss values for each condition. Kernel size 5 consistently minimizes the loss in the EB1-GFP channel, while kernel size 7 performs better in the H2B-RFP channel. During the trials, certain kernel sizes (e.g., 3) led to slower or unstable convergence because the generated feature maps were too noisy or lacked sufficient contextual information to be useful for the model. The feature maps had poor reproducibility under these conditions.

These results underscore the critical role of preprocessing and kernel size optimization in achieving effective model training, demonstrating how M-VEST's adaptability enables precise feature extraction across diverse biological structures. Advanced preprocessing that removes background interference and selects an appropriate kernel size significantly enhances the model's ability to reduce the loss and converge during training. The optimal kernel size for reducing the loss depends on the type of image being analyzed and the preprocessing method used. For example, smaller kernel sizes are more suitable for detecting scattered EB1-GFP signals, whereas larger kernel sizes are necessary to capture the thicker, more contiguous H2B-RFP signals.

**Quality evaluation of ML-derived feature maps**

In addition to the learning curve comparison, the quality of the ML-derived feature maps was evaluated using the key metrics of contrast and entropy, which assess how well the feature maps capture meaningful information (Fig. S3). Contrast refers to the difference in pixel intensity, affecting how clearly distinct structures can be seen. Higher contrast means bright and dark areas are more distinguishable, making key features easier to identify. Entropy measures the complexity or information content in a feature map. Higher entropy suggests more variability in pixel intensities, which can indicate the presence of detailed structures, but may also reflect higher levels of noise in the image. The contrast-to-entropy ratio provides a balance between clarity (contrast) and information richness (entropy) in the feature maps. A higher ratio suggests a good balance in which important features stand out without excessive noise or loss of detail.

The EB1-GFP channel (A1 metaphase vs A1 anaphase) and H2B-RFP channel (A1 metaphase vs AURKA metaphase) were analyzed across kernel sizes of 5, 7, 9, and 11. Kernel size 3 was excluded for the EB1-GFP channel because the learning curve did not reach sufficiently small values or converge, even with the normalization method or the mask + background threshold method (Fig. S2b). For the EB1-GFP channel, both contrast and entropy increased as the kernel size increased from 5 to 11. However, the contrast-to-entropy ratio decreased at larger kernel sizes. This suggests that, although the contrast improved, the feature maps might have become less distinct because of their increased complexity, noise, or even a loss of resolution in fine structures—likely caused by the larger kernel size.

In the H2B-RFP channel, the contrast peaked with a kernel size of 7, whereas the entropy significantly increased with a kernel size of 11. The maximum contrast-to-entropy ratio was achieved with a kernel size of 7, suggesting that this kernel size provides the best balance between contrast, entropy, and resolution for visualizing chromosomal structures in this channel.

These evaluations demonstrate that selecting the right kernel size is crucial for optimizing clarity, feature detection, and resolution in the feature maps. Larger kernel sizes (e.g., 9×9×9 and above) can aggregate information over a broader spatial region, making them more suitable for detecting larger structures. However, while larger kernel sizes may improve contrast, they can also blur finer details and increase noise, ultimately compromising resolution and reducing the precision of feature detection, particularly in smaller structures. These findings align with the earlier conclusion that kernel sizes of 5 and 7 are optimal for the EB1-GFP and H2B-RFP channels, respectively, during model training.

**Considerations on microtubule GTP cap length in different cellular states and its impact on ML results**

In this study, we attempted to extract features from images of GFP-fused EB1, a protein that binds to the growing ends of microtubules (1), a key structure responsible for cell division. EB1 specifically localizes in a comet-like pattern at the growing ends of microtubules, and GFP-EB1 is used as a marker for growing microtubule ends (2, 3). The length of EB1-GFP comets is proportional to the microtubule growth rate (2, 4). Using M-VEST, analysis of the EB1-GFP shapes under different conditions showed that the elongated shape along the long axis is characteristic of fast-growing microtubules, typical of metaphase, which aligns with expectations.

On the other hand, the wider shape along the short axis was suggested by M-VEST analysis to be a feature of anaphase cells or metaphase cells overexpressing AURKA. The thickness of microtubules bound by EB1 does not change significantly under different conditions, so the broader detection of fluorescence intensity suggests an increased EB1-

GFP density. This is consistent with the brightness distribution observed in the images used for ML, confirming that EB1-GFP comets in anaphase or metaphase cells overexpressing AURKA have a higher molecular density. However, this was an unexpected phenomenon that had not been recognized before, prompting us to explore its possible explanations.

This phenomenon could be explained by two primary factors: (1) differences in EB1 locliaztion due to changes in microtubule states during mitosis, or (2) characteristics of EB1-GFP as a marker under the experimental conditions. Both factors likely play a role, and the existence of a mitosis-specific microtubule end structure would be highly intriguing; however, since the latter factor appears to explain the majority of the observed phenomena, we focus our discussion on it below.

Microtubules are dynamic cytoskeletal polymers with a diameter of 25 nm. Their behavior is characterized by “dynamic instability”, a process in which microtubules randomly switch between phases of growth and shrinkage (5). Microtubules polymerize via the net addition of GTP-tubulin subunits at the microtubule plus end, where these subunits subsequently hydrolyze to GDP-tubulin in the microtubule lattice (6). Relatively stable GTP-tubulin subunits form a “GTP cap” at the growing microtubule plus end, which suppresses the switch from growth to shrinkage (6). EB1 recognizes the nucleotide state of tubulin, acting as a proxy for the GTP cap (4).

The length of the GTP cap, as indicated by the EB1-GFP comet length, is generally proportional to the microtubule growth rate (4, 7-9). Cassidy et al. reported that the growth rate-dependent GTP cap lengths in the range of 50–250 nm/s are comparable in vitro and in interphase porcine kidney epithelial cells (LLC-PK1) (8). In our previous work, we reported that average microtubule growth rates in HeLa cells are 0.449 μm/s during metaphase and 0.346 μm/s during anaphase (10) (Fig. S4a, b). Based on the measurements reported by Cassidy et al. (8), the average GTP cap sizes were calculated to be 1132.50 nm during metaphase and 875.00 nm during anaphase (Fig. S4b). However, the EB1-GFP comet length observed in our lattice light-sheet microscopy (LLSM) images, 644.72 nm during metaphase and 501.38 nm during anaphase (Fig. S4e), was shorter than the predicted value, even when measuring the group of the longest comets in either phase. Because the microtubule growth rate varied significantly across different regions of the spindle (Fig. S4a, b), by measuring the comets of microtubules with the fastest growth rate under each condition, it became easier to make comparisons under similar conditions. These comet lengths, based on the fastest-growing microtubules, are approximately 57% of the predicted values during mitosis.

When examining images containing both interphase and mitotic cells, there is a clear tendency for the comets in interphase cells to be longer (Fig. S5a). In interphase cells, the microtubule growth rate is relatively uniform across the entire cell, with a mean of 594.9 ± 7.3 nm/s (n=99), where the value represents the mean ± standard error (SE). The comet length was calculated to be 1270.21 nm (Fig. S5e). Given the predicted GTP cap length of 1497.50 nm based on this growth rate, the comet length is approximately 85% of the predicted value. The expression level of EB1-GFP in these cells is about 40% of that of endogenous EB1(11), indicating a lower ratio of EB1-GFP compared to the endogenous form; therefore, a shorter comet length can be expected. On the other hand, during mitosis, the microtubule growth rate varies depending on the region of the spindle, with the growth rate of astral microtubules extending from the centrosomes being faster (Fig. S4a, b). The variation in growth rate is large, and some astral microtubules extend at speeds greater than 0.6 µm/s. However, as seen in the images in Fig. S5a, even astral microtubules are clearly shorter than the comets in interphase cells.

A plausible explanation for the shorter EB1-GFP comets during mitosis, compared to interphase, is that the increase in the number of microtubules leads to an insufficient supply of EB1-GFP molecules. During mitosis, the number of microtubules increases (12), which can overwhelm the available pool of EB1-GFP molecules, preventing full saturation at the microtubule tips. As a result, the comet lengths are shorter in mitotic cells compared to interphase cells. In this study, additionally, M-VEST indicated that the EB1-GFP density is higher in anaphase than in metaphase. This can be explained by the combined effects of an EB1-GFP shortage, an increase in microtubule number, and a decrease in elongation rate. Our previous measurements showed that the number of EB1-GFP comets in A1 anaphase was approximately three times that of A1 metaphase, with growth rates of 0.346 µm/s in anaphase and 0.449 µm/s in metaphase, respectively (10). Although the slower growth allows EB1-GFP to remain at the tips for longer periods, the overall availability of EB1-GFP molecules decreases further due to the increased number of microtubules, limiting their distribution at the microtubule tips. As a result, the comet length becomes shorter than in metaphase, while EB1-GFP appears more concentrated at the microtubule plus ends due to the reduced growth rate, enabling higher local density despite the overall shortage.

Experimentally, when AURKA was overexpressed in metaphase cells, the number of EB1-GFP comets increased (13), which logically suggested an increase in the number of microtubules. At this time, even in metaphase, the EB1-GFP comet image width was wider, suggesting an increase in EB1-GFP density. Under these experimental conditions, where the EB1-GFP expression level is a limiting factor, the observed higher EB1-GFP densities also support the notion that the EB1-GFP expression level plays a critical role in determining comet density. Additionally, it has been reported that faster microtubule growth rates lead to more dynamic EB1 binding, with EB1 dissociating from GDP-tubulin sites at a faster rate (14). This dynamic behavior causes EB1 to scatter over a wider range, reducing the density of EB1-GFP at any given point along the microtubule. Conversely, slower growth rates allow EB1-GFP to remain concentrated at the microtubule plus ends for longer periods. Together, these opposing effects—EB1-GFP scattering during rapid growth and accumulation during slower growth—likely amplify the differences in EB1-GFP density observed between microtubules with varying growth rates. This highlights how differences in microtubule growth speed contribute significantly to the observed EB1-GFP density variations.

It is still possible that GTP cap formation and EB1 binding might be regulated differently during mitosis and interphase, with potential differences in the stability or extent of GTP cap formation at the microtubule plus ends. During mitosis, the dynamic regulation of microtubules may require faster adjustments to microtubule behavior, which could explain the observed differences in EB1-GFP density between interphase and mitotic cells. Additionally, other factors such as microtubule-associated proteins and the potential for distinct regulation of GTP cap dynamics may also contribute to these differences, and should be considered in future studies to gain a deeper understanding of microtubule dynamics in these cellular states. These factors may also partially influence the behavior of the EB1-GFP marker, leaving open the possibility that mitosis-specific regulatory mechanisms are at play. Exploring such mitosis-specific control mechanisms represents an intriguing avenue for future research.

**Evaluation of M-VEST results and performance**

Based on the above considerations, M-VEST was able to capture subtle changes that cannot be recognized by humans, including EB1-GFP density variations that depend on the microtubule elongation rate. Previous measurements of EB1-GFP comet numbers showed that A1 anaphase had approximately three times as many microtubule growing ends as A1 metaphase (10), and AURKA metaphase had about 1.4 times as many (13). These results correlate well with the current measurements of EB1-GFP comet image width (density) on microtubules. As shown in Fig. 2b, the EB1-GFP short-axis intensity y was approximately 1.7 times higher in A1 anaphase and 1.2 times higher in AURKA metaphase compared with A1 metaphase. The average total widths were estimated as 236 nm for A1 metaphase, 387 nm for A1 anaphase, and 316 nm for AURKA metaphase (Fig. S4c). That is, the A1 anaphase and AURKA metaphase were found to be 151 nm and 80 nm wider than the A1 metaphase. Given that 1 pixel corresponds to 100 nm, this is a subpixel-level difference.

Individual EB1-GFP comets exhibit a broader range of width variations than the average value. These individual differences may enhance the ML model’s ability to detect subtle structural variations with precision close to a single pixel, as the model generalizes from a spectrum of comet widths. By incorporating the unique features of individual comets, the model can capture a more comprehensive structural profile, thereby increasing its sensitivity and accuracy in distinguishing between cellular states. This approach is similar to recent findings in ML-based detection of mitotic cells in pathological specimens, where sensitivity and accuracy are improved by accounting for data diversity and capturing fine variations (15). In conclusion, while individual variations in EB1-GFP comet widths contribute to the model's precision, the broader range of these variations suggests that M-VEST achieves an effective resolution at approximately the single-pixel level, rather than consistently reaching subpixel accuracy.

While this sensitivity is likely influenced by factors such as image patterns, signal-to-noise ratio, and noise levels, the method’s ability to detect differences in chromosomal structures visualized by H2B-RFP—structures that differ significantly from the fine, scattered patterns of EB1-GFP—suggests that M-VEST may perform well across a variety of image types. The high detection capabilities demonstrated in this study indicate that further improvements to the method could enable even finer distinctions in the future.

Despite fewer data points, the learning curve for A1 metaphase vs A1 anaphase (8 trials) dropped to a lower value in the A1 metaphase vs AURKA metaphase case (10 trials), possibly reflecting larger structural differences between metaphase and anaphase. In some combinations, the learning curve did not decrease at all, likely because there were no significant differences between groups, or the differences exceeded M-VEST’s current detection limit. These findings highlight M-VEST’s robustness and remarkable sensitivity, with the ability to detect subpixel-level structural features previously unnoticed by human observation. With further refinement, M-VEST has the potential to reveal even more subtle biological changes.

#### Challenges and potential improvements to M-VEST

M-VEST demonstrates notable sensitivity and adaptability, yet certain technical challenges remain that limit its broader applicability. For instance, the reliance on small kernel sizes restricts the method's ability to detect global structural changes across entire cells or images. This limitation is primarily due to computational resource constraints, as larger kernel sizes significantly increase processing time and memory usage. Future advancements in computation power could allow for the incorporation of larger kernels, enabling more comprehensive analyses without compromising accuracy.

The interpretability of M-VEST-generated feature maps also requires refinement. Developing advanced postprocessing techniques or intuitive visualization tools would make the outputs more accessible to researchers, particularly those outside computational biology fields. For example, overlaying key features with biologically meaningful annotations or offering modular visualization options could enhance usability and facilitate the broader adoption of M-VEST in diverse disciplines.

A current limitation of M-VEST is its single-channel processing capability, which restricts simultaneous analysis of multidimensional datasets. Expanding the framework to integrate multiple fluorescence channels could provide a more holistic understanding of cellular dynamics. This enhancement would be particularly valuable for studying complex biological systems, such as protein-protein interactions or cross-compartmental structural changes in 4D imaging datasets.

Preprocessing methods, including brightness normalization and thresholding, remain critical for ensuring high-quality inputs. While these methods help balance computational efficiency with accuracy, future work could integrate deeper neural networks optimized for complex biological image analysis. These architectures have shown promise in 3D imaging and other fields, suggesting they could further improve M-VEST's capacity to detect intricate structural patterns.

The scalability of M-VEST offers exciting potential for adaptation to various research and clinical contexts. In cancer research, for example, M-VEST could analyze chromosomal abnormalities or microtubule dynamics to provide insights into disease progression and treatment monitoring. Its ability to detect subtle structural changes might also extend to neurodegenerative disorders, where identifying early-stage cellular abnormalities is critical. By expanding its functionality and improving its computational framework, M-VEST could become an indispensable tool for advancing both basic and translational research.

#### Supplementary Materials and Methods

##### Cell culture

The parental HeLa cell line was authenticated by Bio-Synthesis Inc. (Cell Culture STR Profiling and Comparison Analysis, Cat. No.: CL1003). All cell lines tested negative for mycoplasma contamination. HeLa cells and HeLa cells expressing EB1-GFP and histone H2B protein fused to red fluorescent protein H2B-tagRFP (clone A1) (16) were maintained as previously described (11, 17). A1 cells were prepared by introducing H2B-tagRFP into HeLa cells expressing EB1-GFP (clone 2F10), with exogenous EB1-GFP expression levels at approximately 40% of endogenous EB1(11). A1 cells expressing myc-AURKA have previously been described (13), and myc-AURKA expression is about four times higher than endogenous AURKA (13). Control HeLa cells expressing myc-AURKA were generated using lentivirus under the same conditions as described in Ref. (13). In this study, we refer to H2B-tagRFP as H2B-RFP for simplicity, clone A1 as “A1”, the A1 cells with AURKA introduced as “A1-AURKA”, the parental HeLa cells as control, and the control HeLa cells with myc-AURKA introduced as “AURKA”. In A1-AURKA cells, myc-AURKA expression is about four times higher than endogenous AURKA (13).

**Image acquisition**

The data were reused from previously reported studies (13, 16). Data were collected in dual-color and single-color images in dithering mode (fast mode) at intervals of 1.510 s and 0.755 s, respectively. The exposure time per slice was 5 ms. The acquired 3D datasets, obtained through sample scanning in x, y, and s (skew) coordinates, were converted into conventional x, y, z coordinates (“deskewed”). Deconvolution operations were applied using experimentally measured PSFs for each emission wavelength before visualization (see Note 2). The voxel size, which depends on the device state, was calculated to be $0.100\times0.100\times0.217\ \mu m^3$ based on actual measurements. Phases in mitosis were identified by the shape of the chromosomes.

#### Image preprocessing

***Software:*** All image processing steps were performed using the Eos software (18) (https://github.com/tacyas/Eos), IMOD (19), ImageJ, and Fiji (20). Detailed command options and parameters for each processing step are provided in Table S1.

***Image conversion and alignment:*** TIFF images were converted to MRC format using Eos to standardize the pixel dimensions. Resampling was conducted to align the pixel spacing in the X, Y, and Z axes using cubic interpolation. Image rotation was performed using Fiji after determining the tilt of the image, referencing the coverslip surface. Following this, the images were rotated with interpolation applied to correct for the tilt. Cropping was then applied to isolate the region of interest based on the determined coordinates for each image.

***Image intensity normalization:*** Brightness normalization was performed on each 3D image stack at each time point, and was consistently applied across all datasets. As described in Table S1, the normalization process set the high-value area (0.001% of the total) to 0.7 and the low-value area (40% of the total) to 0.15, ultimately normalizing the intensity values to a range of 0.15–0.7. This approach ensures that the brightness values are effectively adjusted to fit within the 0–1 range, minimizing the influence of outliers caused by noise while preventing high-intensity pixels from saturating. Cases in which only normalization was applied are referred to as “normalization” in the paper.

In optimizing the ML conditions, we first used images processed with normalization. However, the learning curve did not decrease satisfactorily, failed to converge, and showed poor reproducibility. Suspecting that the cause might be the brightness in the background of the target structures, we decided to apply advanced processing to remove the background brightness. To achieve this, we first applied a mask to the extracellular regions, and then set a threshold to exclude the background brightness.

***Mask creation and filtering:*** A binary mask was created using the normalized ch0 data (EB1-GFP) to differentiate the areas of interest. Background noise was removed, and regions with a volume of less than half the total cell volume were excluded. A low-pass Gaussian filter was applied to the mask to smooth the transitions, further refining the mask boundaries.

***Mask application and image adjustments:*** The generated mask was applied to both ch0 and ch1 images, and thresholding was conducted to refine the segmentation. Values below a certain threshold were set to zero, while values above the threshold were retained. To adjust the brightness after applying the mask and to exclude the diffuse brightness present in the cytoplasm, the following steps were taken: (1) The average brightness of the regions outside the cell was calculated, and this average brightness was multiplied by −1 (to introduce a negative value). (2) A value of 0.07 was subtracted to further decrease the brightness. This adjustment was intended to minimize the brightness difference at the edges of the mask, reducing the influence of the mask's shape. Specifically, the value added to the image was calculated as (−lower 40% average brightness+6SD)−0.07. As a result, areas in the cytoplasm that originally had low brightness could be assigned brightness values of zero or even negative values. However, after masking, a threshold was set such that values below zero were set to zero, while values above the threshold remained unchanged. The final images were then converted back to TIFF format for further analysis. This process, involving normalization followed by masking and background thresholding, is referred to as "mask + background threshold" in the paper.

**ML tools for image feature extraction**

In this study, we developed an adaptive and interactive ML-based method using a CNN with a single convolutional layer to extract structural feature differences between two groups of 3D image data. The M-VEST code was developed by Apprhythm Inc. (Osaka, Japan) and AISoftware Inc. (Japan). Although the tool is capable of handling 4D datasets (3D space and time), the analysis was limited to 3D in this study. The following sections outline the preprocessing, network architecture, training process, and loss calculation. A detailed workflow, including data preprocessing, augmentation, convolution, and loss calculation, is illustrated in Fig. S2. Parameters specific to each step are listed in Table S2.

***Data preparation and trial design for ML:*** We prepared the following datasets, consisting of EB1-GFP and H2B-RFP channels, for ML: A1 metaphase (11 datasets total; 8 datasets for EB1-GFP after excluding 3 datasets because of strong bleed-through from the H2B-RFP channel; all 11 datasets used for H2B), A1 anaphase (6 datasets), and AURKA metaphase (10 datasets). For each trial, validation data were sequentially selected from the two groups, and the remaining data were used for training. In cases where the number of datasets differed between groups, we reused datasets (starting from the first dataset) in the smaller group, ensuring that the number of trials matched that corresponding to the larger dataset. This resulted in 8 trials for A1 metaphase vs A1 anaphase of the EB1-GFP channel, 10 trials for A1 metaphase vs AURKA metaphase of the EB1-GFP channel, and 11 trials for A1 anaphase vs AURKA metaphase of the H2B-RFP channel. The specific combinations of data and conditions used for the trials are provided in Table S2, where key parameters such as learning rates and batch sizes are also listed.

***Computational resources and program execution:*** The program was executed on the ABCI computational node rt_AF, part of the AI Bridging Cloud Infrastructure (ABCI), which is operated by the National Institute of Advanced Industrial Science and Technology (AIST) (https://abci.ai/). The computational complexity of the 3D convolutional operations made it necessary to modify an existing 3D convolution function to run efficiently on the ABCI system. The modified version of the 3D convolution function is based on the implementation found at https://github.com/funkey/conv4d.

***Image preprocessing and data augmentation:*** The data consisted of two-channel images: EB1-GFP and H2B-RFP channels, each with a size of 256×256×100 pixels (where each image stack contains 100 slices of 256×256 pixels), spanning 30 time points. This resulted in a total of 3000 slices per channel, or 76,800,000 pixels in total for each channel across all time points. The input, however, was processed one channel at a time. The voxel size was set at 0.1×0.1×0.1 μm. The image intensity values were stored in 32-bit floating-point precision to preserve numerical accuracy during processing. The data size for a single time point was approximately 26 MB per channel, resulting in a total of approximately 780 MB per channel across 30 time points.

Brightness correction was not performed as part of the image preprocessing in this tool, as brightness correction and other preprocessing steps had already been completed, as described in the previous section. To ensure that no brightness adjustment occurred, $\alpha$ was set to 0. Data augmentation involving flips and rotation was performed to improve the robustness of the model, resulting in six augmented datasets. Detailed augmentation strategies, such as the specific flip directions and rotations, are illustrated in Fig. S1.

***CNN architecture:*** The convolutional network featured a single 3D convolutional layer, with a filter kernel size ranging from 3×3×3 to 11×11×11 to screen the most suitable size for capturing features that differentiate the two groups. The input 3D image stacks $X$, with dimensions of $(x,y,z)$, were processed through the convolutional layer using a learned weight parameter matrix $W_{pq}$. The convolution operation was followed by pooling to downsample the feature maps, and the output was passed through a nonlinear activation function to generate feature maps representing specific aspects of the input data. M-VEST learns the optimal weights for each kernel during the training process. These weights are iteratively refined using regularization techniques, such as the Frobenius norm, to control their magnitude and ensure effective feature extraction.

***Training process:*** First, the dataset was split into mini-batches containing one training example. The Adam optimizer (21) with an initial learning rate of 0.001 was then applied, and the model was trained for 10–20 epochs, depending on the dataset and specific filter kernel size. The training process was designed to iteratively minimize the loss function based on the Frobenius norm, which penalized deviations of the weight matrices from the identity matrix. The learned weights were further regularized using the L1 norm to prevent overfitting and ensure that the CNN filters maintained a similar scale between the input image and the output feature map.

***Output (feature map generation):*** Once the training process reached the set number of epochs, feature maps were generated by applying convolutional filters to the test datasets.

Each filter was optimized during training to detect specific patterns or features, such as edges, textures, or more complex structures, within the images. These filters assigned different levels of importance to regions of the image, emphasizing relevant features and suppressing less-significant areas.

***Model validation and tuning of the learning process:*** To optimize the weight parameters of the convolutional filters, we used a loss function that incorporated both label prediction and a Frobenius norm regularization term to penalize deviations. The probability function *PF* was calculated using the output feature map $F(X / W_{pq})$. If *PF* exceeded 0.5, the example was classified as positive; otherwise, it was classified as negative. The Frobenius norm coefficient λ was set to 2,000,000 to strongly regularize the learned filters, ensuring the CNN filter weights $W_{pq}$ remained balanced. The L1 norm was applied to ensure that the gain of the CNN filters, represented by the weight matrix $W_{pq}$, remained at approximately 1, keeping the pixel values of the input image and the output feature map similarly scaled. The overall loss function was defined as follows:

$$\widehat{W}_{pq} = \underset{W_{pq}}{\operatorname{argmin}} \left[ \sum_{i=1}^{N} \left\{ \frac{1}{N_n}(1 - l_i) \cdot \hat{l}_i - \frac{1}{N_p} l_i \cdot \hat{l}_i \right\} + \lambda \left| \sqrt{\sum_{w_{pq} \in W_{pq}} w_{pq}^2} - 1 \right| \right]$$

where:

- $l_i$ is the label for each example, set to 1 if the example is positive and 0 if it is negative;
- $\hat{l}_i$ is the predicted label for example i, set to 1 for positive and 0 for negative predictions;
- $N_p$ and $N_n$ represent the total number of positive and negative examples, respectively;
- $W_{pq}$ is the convolutional filter weight matrix applied to the input data;

- $w_{pq}$ are the elements of the weight matrix $W_{pq}$, and the summation $\sum w_{pq} \in W_{pq}$ is taken over all elements of $W_{pq}$;
- $\lambda$ is the regularization coefficient controlling the strength of the Frobenius norm penalty, ensuring that the weights remain balanced.

The regularization term ensured that the filter weights remained balanced, preventing overfitting. The loss function was minimized using the Adam optimizer, which adjusted the model weights iteratively through backpropagation, enabling effective convergence during training. Model performance was validated on a test dataset, with hyperparameters such as the regularization coefficient λ and epoch size adjusted to optimize convergence.

***Visualization of 3D feature maps:*** Once the model demonstrated satisfactory performance, the generated feature maps were evaluated by humans. To visualize the 3D feature maps, we used ImageJ to open the images and observe the 3D maps as slice images at specific time points. Additionally, we visualized the overall distribution of signal intensity as a movie by viewing either a particular slice plane or using the average intensity projection method. The images were displayed using a fire lookup table (LUT) to enhance signal visibility, with brightness adjustments applied to make the signals easier to observe. A calibration bar indicated the displayed brightness range.

In these visualized feature maps, the background appeared in shades ranging from black to white, reflecting the varying activation values produced by the convolutional filters in response to different input patterns. This variation in background intensity results from the model's ability to capture features through both positive and negative activation values, with backgrounds sometimes appearing completely black or white. These variations reflect feature intensity changes detected by the network and do not interfere with the interpretation of the maps.

During CNN training, the model learned to classify input images by identifying relative feature strengths, with pixel values in the feature maps representing how strongly a feature contributes to classification. The absolute sign of these values—positive or negative—was arbitrary and did not impact the model's performance. It is the relative contrast across the feature map that is important because classification depends on the pattern of intensities rather than their absolute values. The model's initial random weight assignments determined whether specific features were labeled as positive or negative, but this did not influence the effectiveness of the CNN's classification.

Overall, the variations in activation values were part of the network's natural learning process, helping it adapt to capture the most informative spatial features. Consequently, variations in background brightness or contrast did not influence the accuracy of feature detection or classification.

**Feature map evaluation using contrast and entropy**

The ML-generated feature maps were evaluated across various kernel sizes (5×5×5, 7×7×7, 9×9×9, and 11×11×11) to identify the optimal size for capturing distinguishing features. Calculations were performed using Python, specifically leveraging the libraries numpy and scipy to compute the key metrics of mean contrast, mean entropy, and contrast-to-entropy ratio. The kernel size 3×3×3 was excluded from further analysis because of the poor performance of its learning curve, which failed to achieve sufficient convergence.

Consequently, it was not considered for detailed feature map evaluations in the subsequent analyses.

***Contrast measurement:*** The mean contrast, representing image sharpness, was calculated using the numpy library by determining the mean and standard deviation of pixel intensity. Higher contrast indicates clearer differentiation between structures. Contrast values were averaged across all feature maps for each kernel size.

***Entropy calculation:*** The entropy, quantifying image complexity or randomness, was calculated using the scipy.stats.entropy function. A pixel intensity histogram was normalized to probabilities, and the Shannon entropy was computed. Higher entropy values suggest greater complexity, indicating either meaningful details or noise.

***Contrast-to-entropy ratio:*** The contrast-to-entropy ratio was computed by dividing the mean contrast by the mean entropy for each feature map. This metric was used to evaluate the trade-off between clarity (contrast) and complexity (entropy). A higher ratio indicates that the feature map preserves detail without introducing excessive noise. The kernel size with the highest contrast-to-entropy ratio was selected for further analysis on the basis that it provided the best balance between image sharpness and relevant detail.

**LLSM image data analysis**

Analysis of the EB1-GFP comet intensity distribution (in both the short- and long-axis directions) was performed using the 2D maximum intensity projection of LLSM 3D image stacks in ImageJ. The signal distribution of EB1-GFP comets on astral microtubules was measured using line plots generated in ImageJ. To measure the width (short axis) of EB1-GFP comet intensities in the images used for ML, we used the mask + background threshold dataset used for ML. The comet length depends on the stage of elongation, so we measured comets with an average shape at the elongation stage. Additionally, because part of the EB1-GFP comet intensity may have been reduced by the mask + background threshold process, we also measured the EB1-GFP comet length from images in the normalized dataset. Note that, in this case, the maximum comet length was determined by selecting longer comets from A1 metaphase and anaphase cells. Line profiles were measured using a 3-pixel-wide line. The calculation of the mean and standard error (SE), plotting, and statistical analysis were conducted using Python. For statistical analysis, a two-sample t-test was conducted at each data point to compare the means between the A1 metaphase group and the other groups. The p-values were computed for each comparison and plotted as $p \leq 0.05$*, $p \leq 0.01$**, and $p \leq 0.001$***.

To determine the short-axis width of the EB1-GFP comets, the following calculations were performed using Python. For each condition, the average intensity across the replicates was calculated. Additionally, SE was computed for each position to represent the variability in the measurements. To reduce noise in the intensity data, we applied a Gaussian smoothing function using a standard deviation (sigma) of 0.5 (a general Gaussian function was not suitable for these specific data as it could not capture the shape of the intensity profiles effectively). This smoothing process preserved the overall structure of the data while mitigating fluctuations. A line plot was generated for each condition, displaying the smoothed intensity data as a function of position (nm). Error bars, representing SE, were added to each data point. The width of the intensity profile at Y = 0.05 and Y = 0.1 was calculated for each condition. For this, we identified the positions on both the rising and falling edges of the intensity curve at which the intensity crossed the specified Y value. The difference between these positions was defined as the width.

To facilitate direct comparison with the EB1 comet tail length results reported by Cassidy et al. (8), the comet tail length was determined based on exponential decay fitting(22, 23) applied to the averaged intensity profiles of metaphase and anaphase data. Intensity profiles were extracted from the aligned dataset, and the data was grouped into "anaphase" and "metaphase" categories. For each group, the mean intensity at each position was calculated across all columns in the dataset to generate averaged intensity profiles, which were then subjected to exponential decay fitting. The fitting was initiated from the first data point immediately following the intensity peak to ensure that only the decay phase of the comet was analyzed. The exponential decay was modeled using the equation $y=a \cdot \exp(-b \cdot x)+c$, where $a$ represents the initial amplitude, $b$ is the decay constant, $c$ is an offset representing the baseline intensity, and $x$ represents the distance in nanometers. The decay constant ($b$) obtained from the fitting was used to calculate the characteristic decay length ($d$) as the inverse of $b$ ($d=1/b$). To ensure compatibility with prior studies, this length ($d$) was scaled by a factor of 1/0.37. This scaling factor was introduced to provide a more comparable representation of the comet tail length and reflects the characteristic distance over which the intensity decreases in the context of earlier work. In this study, the term "characteristic decay length" refers to the direct calculation based on the exponential decay constant, while the term "scaled comet tail length ($d_{\text{scaled}}$)" represents the adjusted value derived by applying the scaling factor to $d$. Both metrics were calculated and visualized to provide a comprehensive analysis of the EB1 comet structure. The results of the exponential decay fitting were overlaid on the averaged intensity profiles for both metaphase and anaphase groups, and the scaled comet tail lengths ($d_{scaled}$) were included in the figure legends to summarize the outcomes of the analysis. All analyses, including data processing, exponential decay fitting, and visualization, were performed using Python. The numerical calculations were carried out using the NumPy library, and all visualizations were generated with Matplotlib. The workflow was designed to facilitate direct comparison with previously published studies on EB1 dynamics (8, 22, 23).

The data on the microtubule growth speed in A1 cells based on spatial statistics were reused from previously published work (10). The data on the number of EB1-GFP comets in A1 metaphase cells and AURKA metaphase cells were also taken from previous research (13). The tracking of EB1-GFP comets in interphase cells was performed using Imaris, as previously reported (10).

To visualize the trajectories of EB1-GFP over time (Fig. 2c), an average intensity projection was generated to obtain the overall distribution of fluorescence using ImageJ. Prior to the creation of the projection, the position of the spindles, which undergo rotational movement within the cell, was corrected using Imaris, based on the centrosome as a reference, as previously reported (10).

**Fluorescent antibody labeling and fluorescence microscopy**

The following commercial primary antibodies were used: anti-acetylated histone2B polyclonal rabbit antibody (Millipore, 07-373, 3092508); anti-acetylated histone 3 polyclonal rabbit antibody (Merck, 06-599, 3260200); anti-acetylated histone 4K16 recombinant rabbit antibody (Abcam, ab109463, GR284778-8). Hoechst 33342 solution

(346-07951; Lot# FN027; Dojin, Japan) was used as a nuclear marker. Immunofluorescent staining of fixed cultured cells was performed as described previously (11, 17, 24). The immunostained specimens were observed using an LSM880 confocal microscope with a Plan-APOCHROMAT 63×/1.4 NA oil immersion for SR objective, four laser lines (405 nm; Multi-Ar, 458, 488, 514 nm; DPSS, 561 nm; He-Ne, 633 nm), and a GaAsP detector (Carl Zeiss). The proportion of acetylated histones was calculated using the method described in Ref. (25), and the ratio between the signal was determined from acetylated histone antibodies and the Hoechst 33342 signal. The analysis was performed using ImageJ, where the chromosome regions were manually outlined, and the intensity from different channels was measured and calculated.

**Statistics**

Statistical evaluation was carried out by the Student's t-test (two sided). P-values and sample sizes are shown in each figure.

**Data and materials availability**

Data are available on request from the authors. Materials used in this study are available for distribution after a materials transfer agreement.

**Code availability**

The image processing software tool Eos, used for image preprocessing, is available at https://github.com/tacyas/Eos. The code for the M-VEST method will be made available for collaborative research purposes until this paper is published. Other computer codes are available on request from the authors.

#### Legend for Tables S1 and S2, and Movies S1 to S6

**Table S1.**

This table provides the detailed preprocessing steps applied to the images used in the ML analysis, including processing method, software and commands, and corresponding input parameters.

**Table S2.**

This table summarizes the experimental setup for M-VEST analysis across various ML combinations. Each sheet contains details for different comparisons. The Trial Combinations section lists the specific cells analyzed for each condition and trial.

**Movie S1.**

A 3D stack from the third time point of the original images and feature maps for the EB1-GFP channel in A1 metaphase vs A1 anaphase training is shown, moving from top to bottom. In the feature maps, the top row highlights pixels that are characteristic of metaphase, while the bottom row highlights pixels that are characteristic of anaphase. The display brightness ranges are 0–0.2 for the EB1-GFP channel and 0–0.5 for the H2B-RFP channel. The feature maps are shown using a fire LUT, with the displayed range indicated by the calibration bar. Slice numbers are displayed in the top left. Scale bar: 5 µm.

**Movie S2.**

A time-lapse movie showing the original images (maximum intensity projection of 50 slices) and feature maps (average intensity projection of 50 slices, fire LUT) for the EB1-GFP channel in A1 metaphase vs A1 anaphase training. In the feature maps, the top row highlights pixels that are characteristic of metaphase, while the bottom row highlights pixels that are characteristic of anaphase. The display brightness ranges are 0–0.2 for the EB1-GFP channel and 0–0.5 for the H2B-RFP channel. The feature maps are shown using a fire LUT, with the displayed range indicated by the calibration bar. Scale bar: 5 µm; time: s:ms.

**Movie S3.**

A 3D stack from the third time point of the original images and feature maps for the EB1-GFP channel in A1 metaphase vs AURKA metaphase training is shown, moving from top to bottom. In the feature maps, the top row highlights pixels that are characteristic of metaphase, while the bottom row highlights pixels that are characteristic of anaphase. The display brightness ranges are 0–0.2 for the EB1-GFP channel and 0–0.5 for the H2B-RFP channel. The feature maps are shown using a fire LUT, with the displayed range indicated by the calibration bar. Slice numbers are displayed in the top left. Scale bar: 5 µm.

**Movie S4.**

A time-lapse movie showing the original images (maximum intensity projection of 50 slices) and feature maps (average intensity projection of 50 slices, fire LUT) for the EB1-

GFP channel in A1 metaphase vs AURKA metaphase training. In the feature maps, the top row highlights pixels that are characteristic of metaphase, while the bottom row highlights pixels that are characteristic of anaphase. The display brightness ranges are 0–0.2 for the EB1-GFP channel and 0–0.5 for the H2B-RFP channel. The feature maps are shown using a fire LUT, with the displayed range indicated by the calibration bar. Scale bar: 5 µm; time: s:ms.

**Movie S5.**

A 3D stack from the third time point of the original images and feature maps for the H2B-RFP channel in A1 metaphase vs AURKA metaphase training is shown, moving from top to bottom. In the feature maps, the top row highlights pixels that are characteristic of metaphase, while the bottom row highlights pixels that are characteristic of anaphase. The display brightness ranges are 0–0.2 for the EB1-GFP channel and 0–0.5 for the H2B-RFP channel. The feature maps are shown using a fire LUT, with the displayed range indicated by the calibration bar. Slice numbers are displayed in the top left. Scale bar: 5 µm.

**Movie S6.**

A time-lapse movie showing the original images (maximum intensity projection of 50 slices) and feature maps (average intensity projection of 50 slices, fire LUT) for the H2B-RFP channel in A1 metaphase vs AURKA anaphase training. In the feature maps, the top row highlights pixels that are characteristic of metaphase, while the bottom row highlights pixels that are characteristic of anaphase. The display brightness ranges are 0–0.2 for the EB1-GFP channel and 0–0.5 for the H2B-RFP channel. The feature maps are shown using a fire LUT, with the displayed range indicated by the calibration bar. Scale bar: 5 µm; time: s:ms.

**Fig. S1 (to be continued)**

#### Learning feature extraction filters from 3D image stacks through solving a Convolutional Neural Network (CNN) learning problem with a single convolutional layer

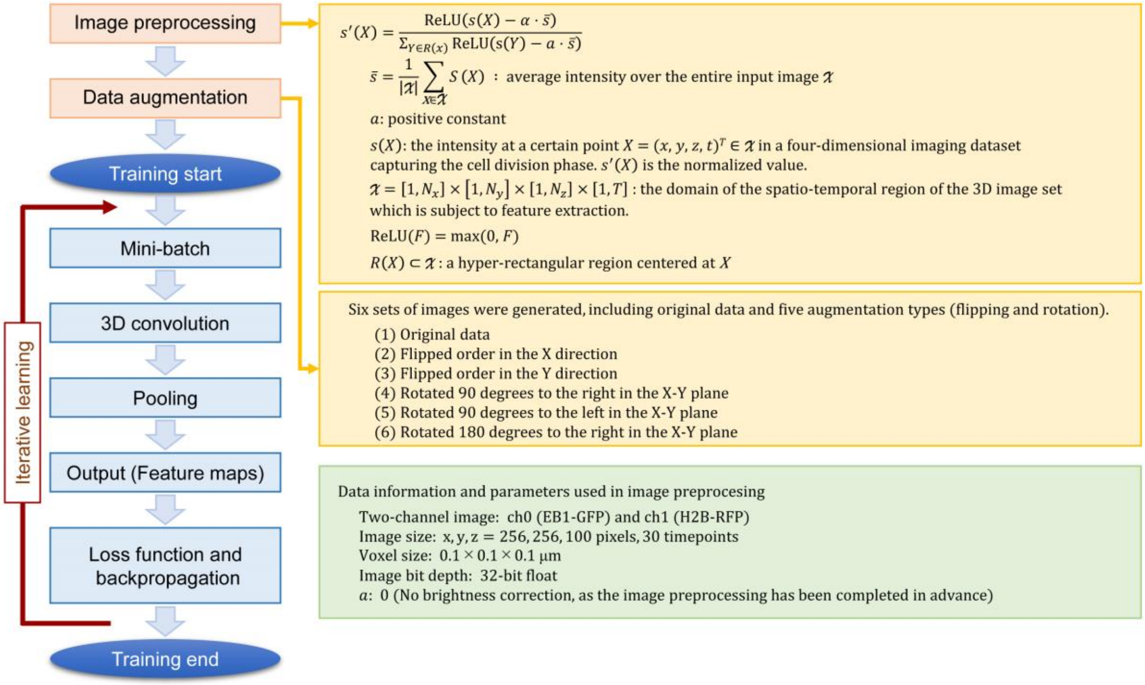

**Fig. S1 (continue)**

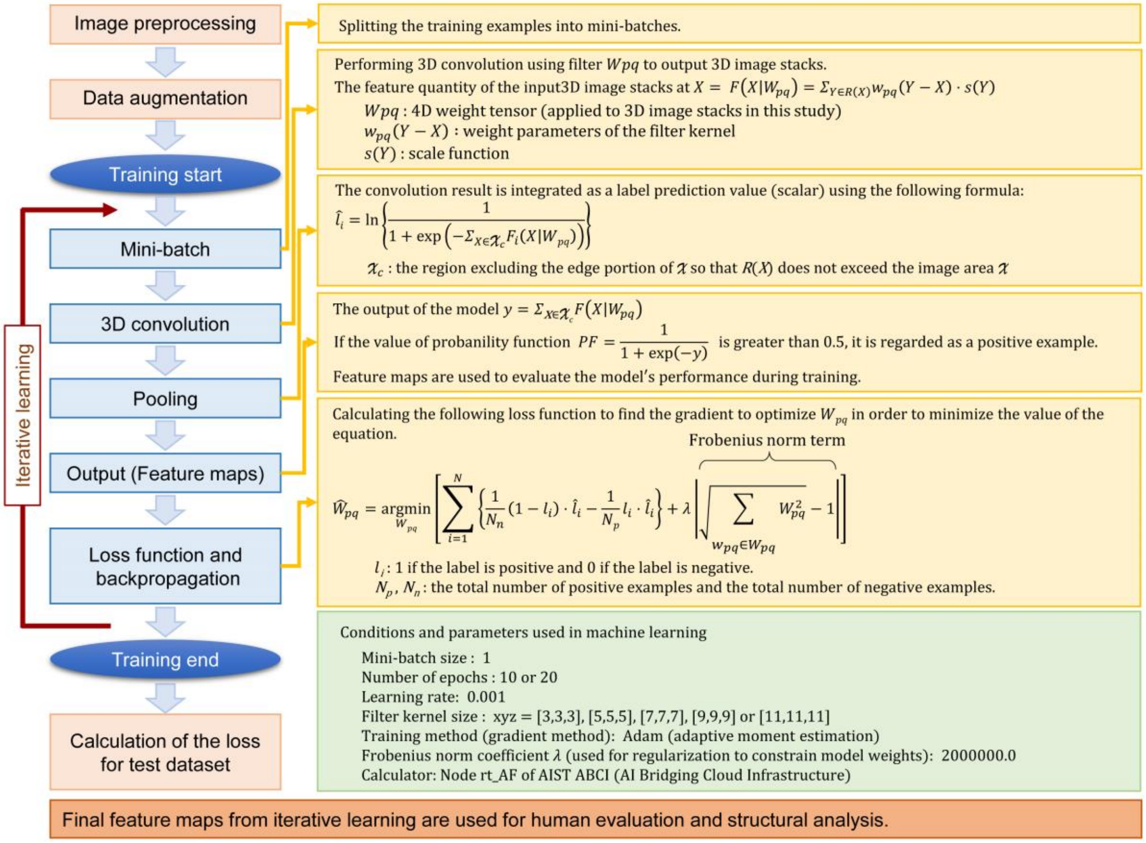


**Fig. S1. Workflow for learning feature extraction filters in M-VEST using a single convolutional layer applied to 3D image stacks.** This workflow, as part of the overall M-VEST process (see Fig. 1 for overall workflow), illustrates the iterative procedure for learning feature extraction filters from 3D image stacks using a CNN with a single convolutional layer. In this method, multiple trials are conducted by selecting different combinations of training and test datasets from the available data, allowing the model to generalize effectively across variations. Key steps include image preprocessing, data augmentation, 3D convolution, and pooling. The output from each trial is analyzed through feature maps, with parameters such as image size, voxel size, and ML conditions (minibatch, learning rate, filter kernel size) provided. The final feature maps generated across trials are then used for human evaluation and structural analysis.

**Fig. S2**

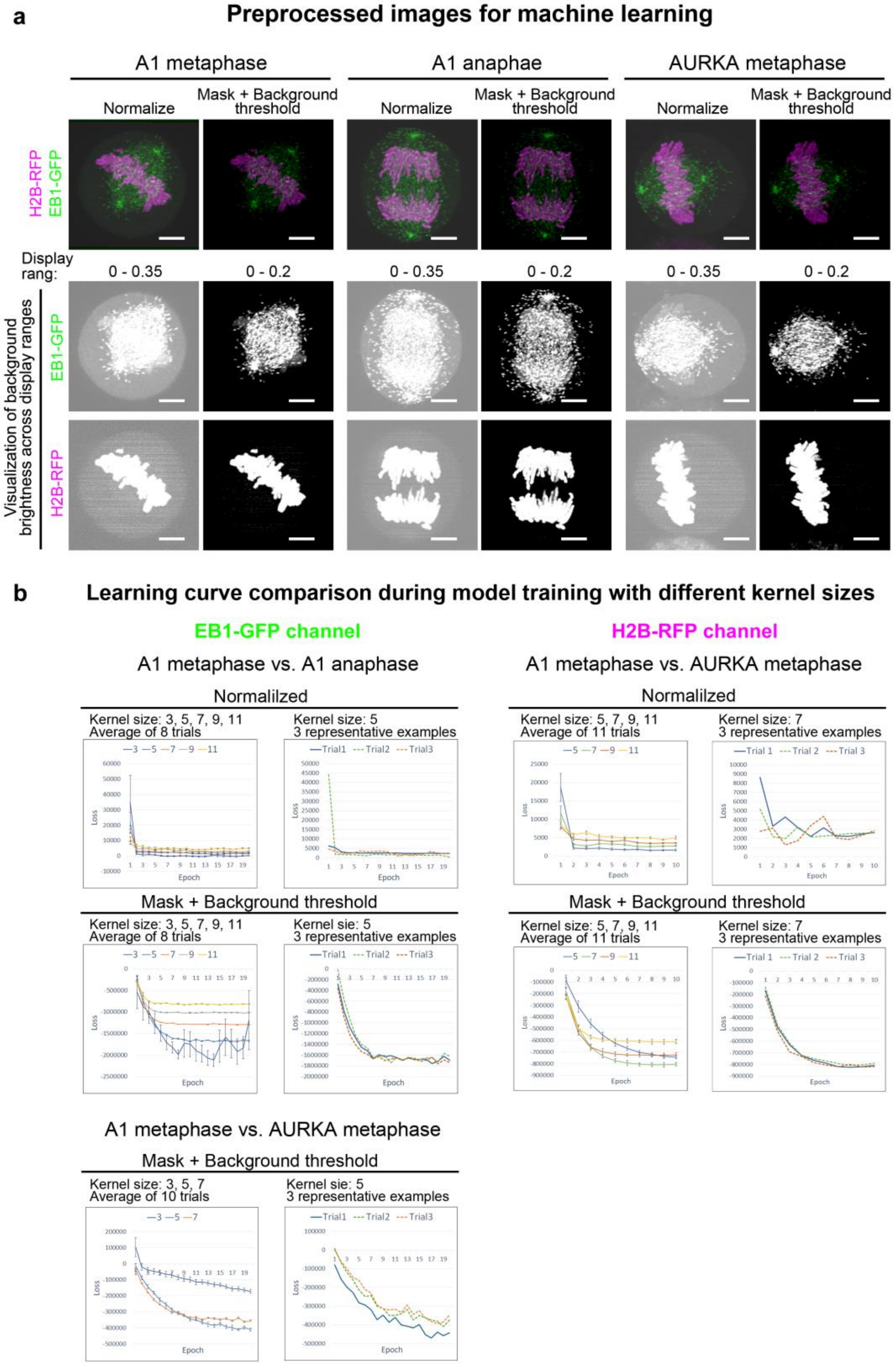
a
Preprocessed images for machine learning
A1 metaphase
A1 anaphae
AURKA metaphase
Normalize
Mask + Background threshold
H2B-RFP
EB1-GFP
Display rang:
0 - 0.35
0 - 0.2
Visualization of background brightness across display ranges
b
Learning curve comparison during model training with different kernel sizes
EB1-GFP channel
H2B-RFP channel
A1 metaphase vs. A1 anaphase
A1 metaphase vs. AURKA metaphase
Normalilzed
Mask + Background threshold
Kernel size: 3, 5, 7, 9, 11
Average of 8 trials
Kernel size: 5
3 representative examples
Kernel sie: 5
Kernel size: 5, 7, 9, 11
Average of 11 trials
Kernel size: 7
Kernel size: 3, 5, 7
Average of 10 trials
Trial1
Trial2
Trial3
Loss
Epoch

**Fig. S2. Preprocessed images and learning curve comparison during model training with different kernel sizes.** (**a**) Preprocessed images for ML. Representative images from the EB1-GFP and H2B-RFP channels in control metaphase, control anaphase, and AURKA metaphase cells, used for ML. The original images in the top row show a brightness range of 0–1. The images were preprocessed with to form normalization and mask + background threshold datasets. Display range variations show how background brightness is visualized across different processing conditions. Scale bars: 5 µm. (**b**) Learning curve comparison during model training with different kernel sizes. Loss curves for the EB1-GFP and H2B-RFP channels are shown for model training under different preprocessing conditions (normalized and mask + background threshold) and kernel sizes (3, 5, 7, 9, 11). Normalization alone did not reduce the loss sufficiently. In the EB1-GFP channel, model training did not progress well with kernel size 3, even with mask + background threshold preprocessing, but other kernel sizes resulted in good convergence despite differences in loss values. The results compare A1 metaphase vs A1 anaphase for EB1-GFP channel and A1 metaphase vs AURKA metaphase for H2B-RFP channel. For the EB1-GFP channel in A1 metaphase vs AURKA metaphase, only kernel sizes 3, 5, and 7 were used based on the results of A1 metaphase vs A1 anaphase. Each graph presents either the average of multiple trials or representative examples.

Fig. S3

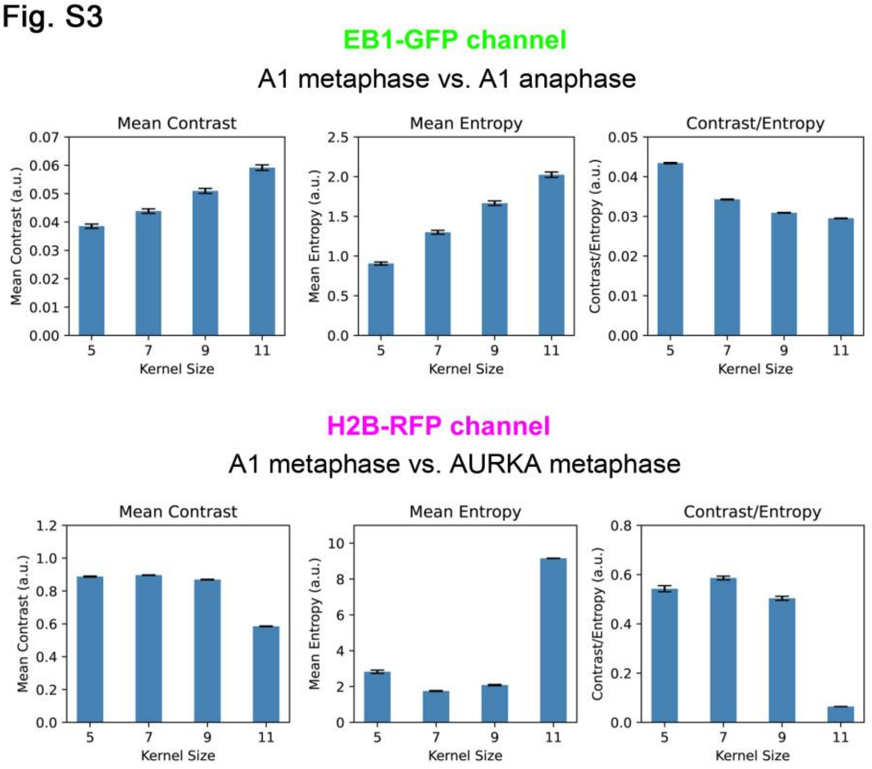


**Fig. S3. Contrast and entropy analysis across different kernel sizes for the EB1-GFP and H2B-RFP channels.** For the EB1-GFP channel (top row), comparisons were made between A1 metaphase and A1 anaphase across kernel sizes 5, 7, 9, and 11. The bar plots show the changes in contrast (left), entropy (middle), and the contrast-to-entropy ratio (right) for each kernel size. For the H2B-RFP channel (bottom row), comparisons were made between A1 metaphase and AURKA metaphase across the same kernel sizes. The bar plots similarly display contrast (left), entropy (middle), and the contrast-to-entropy ratio (right). Error bars represent the standard error (SE).

**Fig. S4**

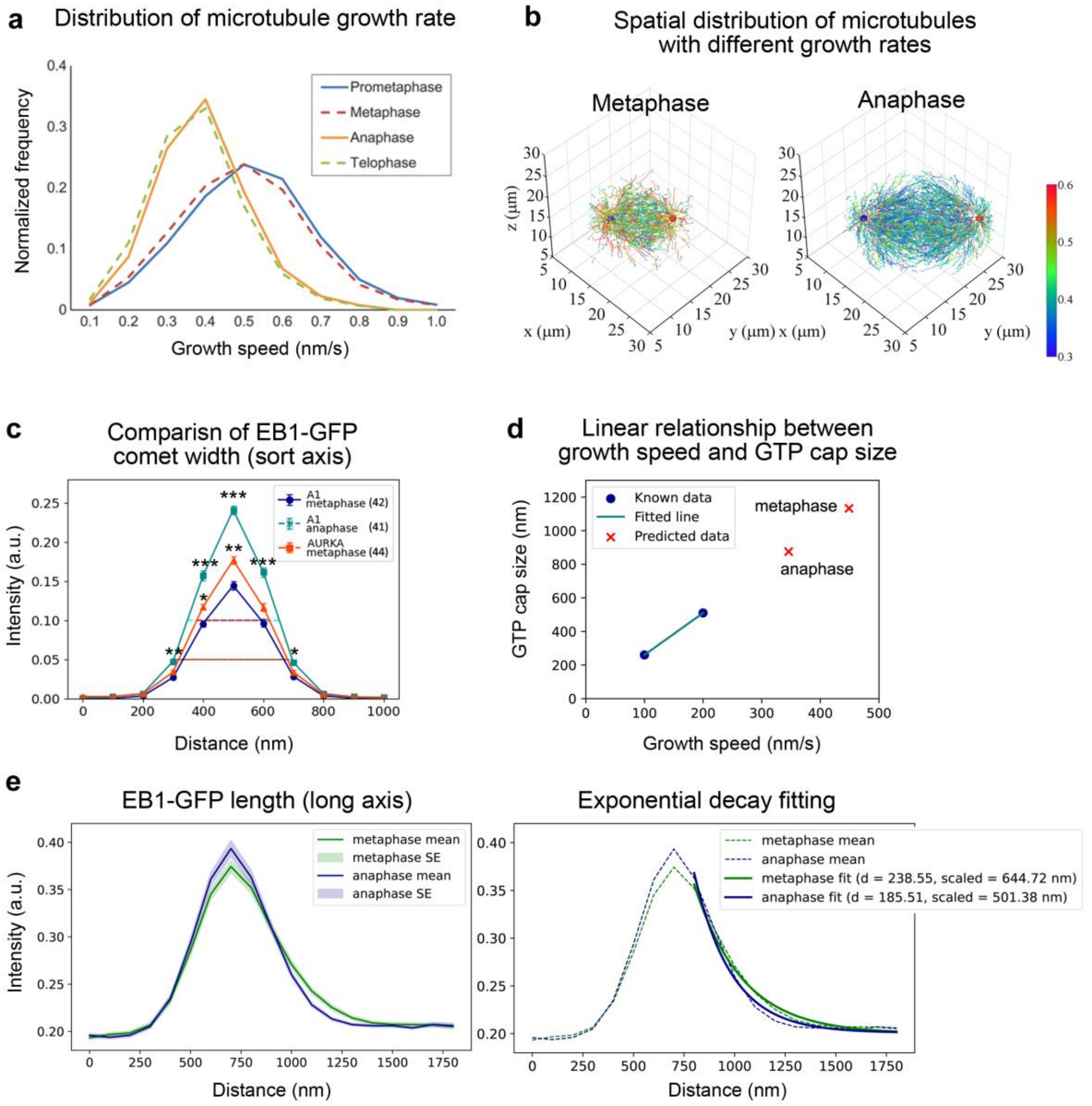


**Fig. S4. Analysis of EB1-GFP comet width, growth speed, and microtubule dynamics across different phases.** (**a**) Frequency distribution of EB1-GFP comet speed for each mitotic phase obtained by averaging data obtained from three cells (8374 and 18964 tracks for metaphase and anaphase, respectively). Data reused from Reference 10. (**b**) Merged trajectories of EB1-GFP comets in metaphase and anaphases. Blue and red triangles indicate centrosomes. Orange and cyan dots indicate the start and end positions of trajectories, respectively. The colored bar indicates the mean speed of EB1-GFP comet trajectories (0.3–0.6 µm/s). Data reused from Reference 13. (**c**) Analysis of EB1-GFP comet width in the short-axis direction. The plot illustrates the intensity profiles of EB1-GFP comets for the A1 metaphase, A1 anaphase, and AURKA metaphase conditions. The profiles, derived from the data shown in Fig. 2b, were smoothed using Gaussian smoothing, and the comet width was analyzed in the short-axis direction. Gaussian smoothing was employed to preprocess the data, because Gaussian fitting could not be directly applied to the intensity profiles due to the limited number of pixels, which made the profiles unsuitable for robust Gaussian fitting. Dashed lines indicate the calculated widths at specific intensity levels of Y = 0.05 and Y = 0.1. At Y = 0.05, the comet widths are 335 nm for A1 metaphase, 394 nm for A1 anaphase, and 362 nm for AURKA metaphase. At Y = 0.1, the widths are 185 nm for A1 metaphase, 306 nm for A1 anaphase, and 241 nm for AURKA metaphase. These results demonstrate that the A1 anaphase condition exhibits the widest comets at both intensity thresholds, highlighting notable differences in comet width between the conditions. The difference between A1 metaphase and AURKA metaphase is 27 nm at Y = 0.05 and 56 nm at Y = 0.1; because one pixel corresponds to 100 nm, M-VEST is able to detect subpixel differences. Error bars represent SE at each data point. $p \leq 0.05$*, $p \leq 0.01$**, and $p \leq 0.001$***. (**d**) Linear relationship between growth speed and GTP cap size. A plot showing the fitted linear relationship between microtubule growth speed (nm/s) and GTP cap size (nm) based on known or previously reported data (8). The GTP cap size was calculated based on the average microtubule growth speeds of 0.449

μm/s in metaphase and 0.346 μm/s in anaphase (10). (**e**) Fluorescence intensity profile of EB1-GFP along the long axis during metaphase (green) and anaphase (blue) (left). Note that EB1-GFP comet length was measured from the longer comets present in each cell. Solid lines represent the mean fluorescence intensity profiles, and shaded regions indicate the standard error (SE). Although metaphase comets tend to be longer, there is no significant difference. On the right, the profiles were fitted with a Bi-Gaussian model to account for the asymmetry of the intensity distribution. Exponential decay fitting was performed starting from the point immediately after the Bi-Gaussian peak, as described by Bieling et al(22). The characteristic comet lengths ($d$) for metaphase and anaphase were calculated from the inverse of the exponential decay constant. These results highlight distinct differences in comet length between metaphase and anaphase.

**Fig. S5**

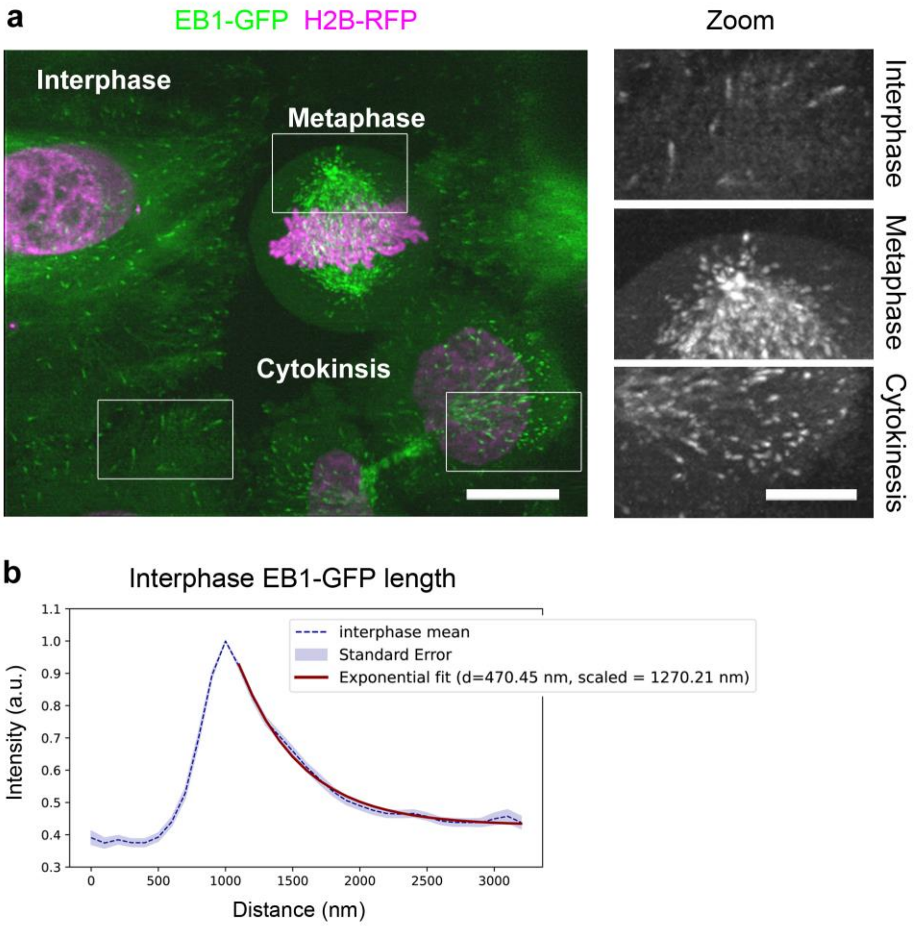


**Fig. S5.** EB1-GFP dynamics across cell cycle stages and quantification of EB1-GFP comet lengths. (**a**) Image showing EB1-GFP and H2B-RFP across different cell cycle stages. Left: representative image showing EB1-GFP (green) and H2B-RFP (magenta) in interphase, metaphase, and cytokinesis. Right: magnified views of the boxed areas in the left panel. Only the EB1-GFP channel is shown in grayscale. EB1-GFP comets in interphase cells are longer than those in mitotic cells, with a microtubule growth rate of 594.9 ± 7.3 nm/s (n=99, instantaneous speed). The predicted GTP cap size is 1497.50 nm (See Fig. S4b). Brightness is not informative because individual brightness correction was not applied for each cell. Scale bars: 10 µm (left), 5 µm (right). (**b**) Quantification of EB1-GFP comet length during interphase. Fluorescence intensity along the length of EB1-GFP comets in interphase cells is shown. The data is normalized, and the curve represents the mean fluorescence intensity profile. The shaded area shows the standard error. The exponential fit to the data is shown in red with the corresponding tail length (d = 470.45 nm, scaled = 1270.21 nm). The fit describes the decay of EB1-GFP comet intensity, starting from the point immediately next to the peak, as described by Bieling et al. (22). Therefore, the actual EB1-GFP comet length is approximately 84% of the predicted value.
