## Supplementary Table S2 for "Comparative Extraction of Cellular Features from High-Resolution Volume Imaging"

| | | | | | | | | | |
|---|---|---|---|---|---|---|---|---|---|
| **Channel:** | EB1-GFP | | | | | | | | |
| **ML combination:** | A1 metaphase vs. A1 anaphase | | | | | | | | |
| **Trial Combinations:** | | Trial 1 | Trial 2 | Trial 3 | Trial 4 | Trial 5 | Trial 6 | Trial 7 | Trial 8 |
| | A1 metaphase | cell 2 | cell 4 | cell 5 | cell 6 | cell 7 | cell 8 | cell 10 | cell 11 |
| | A1 anaphase | cell 1 | cell 2 | cell 3 | cell 4 | cell 5 | cell 6 | cell 1 | cell 2 |
| **Augmentation:** | default | | | | | | | | |
| | flip_horizontal | | | | | | | | |
| | flip_vertical | | | | | | | | |
| | rot90_right | | | | | | | | |
| | rot90_left | | | | | | | | |
| | rot180 | | | | | | | | |
| **Batch size** | 1 | | | | | | | | |
| **Learning_Rate:** | 0.001 | | | | | | | | |
| **Epoch:** | 20 | | | | | | | | |
| **Frobenius norm ($\lambda$):** | 2000000 | | | | | | | | |

**Table S2**

| | | | | | | | | | | | |
|---|---|---|---|---|---|---|---|---|---|---|---|
| **Channel:** | EB1-GFP | | | | | | | | | | |
| **ML combination:** | A1 metaphase vs. AURKA metaphase | | | | | | | | | | |
| **Trial Combinations:** | | Trial 1 | Trial 2 | Trial 3 | Trial 4 | Trial 5 | Trial 6 | Trial 7 | Trial 8 | Trial 9 | Trial 10 |
| | A1 metaphase | cell 2 | cell 4 | cell 5 | cell 6 | cell 7 | cell 8 | cell 10 | cell 11 | cell 2 | cell 4 |
| | AURKA metapha: | cell 1 | cell 2 | cell 3 | cell 4 | cell 5 | cell 6 | cell 7 | cell 8 | cell 9 | cell 10 |
| **Augmentation:** | default | | | | | | | | | | |
| | flip_horizontal | | | | | | | | | | |
| | flip_vertical | | | | | | | | | | |
| | rot90_right | | | | | | | | | | |
| | rot90_left | | | | | | | | | | |
| | rot180 | | | | | | | | | | |
| **Batch size** | 1 | | | | | | | | | | |
| **Learning_Rate:** | 0.001 | | | | | | | | | | |
| **Epoch:** | 20 | | | | | | | | | | |
| **Frobenius norm ($\lambda$):** | 2000000 | | | | | | | | | | |

**Table S2**

| | |
|---|---|
| **Channel:** | H2B-RFP |
| **ML combination:** | A1 metaphase vs. AURKA metaphase |

**Trial Combinations:**

| | Trial 1 | Trial 2 | Trial 3 | Trial 4 | Trial 5 | Trial 6 | Trial 7 | Trial 8 | Trial 9 | Trial 10 | Trial 11 |
|---|---|---|---|---|---|---|---|---|---|---|---|
| A1 metaphase | cell 1 | cell 2 | cell 3 | cell 4 | cell 5 | cell 6 | cell 7 | cell 8 | cell 9 | cell 10 | cell 11 |
| AURKA metapha: | cell 1 | cell 2 | cell 3 | cell 4 | cell 5 | cell 6 | cell 7 | cell 8 | cell 9 | cell 10 | cell 1 |

| | |
|---|---|
| **Augmentation:** | default |
| | flip_horizontal |
| | flip_vertical |
| | rot90_right |
| | rot90_left |
| | rot180 |
| **Batch size** | 1 |
| **Learning_Rate:** | 0.001 |
| **Epoch:** | 10 |
| **Frobenius norm (λ):** | 2000000 |
