## Supplementary Table S1 for "Comparative Extraction of Cellular Features from High-Resolution Volume Imaging"

| Purpose | Processing | | Software | Eos command | Input Parameters | | Option |
|---|---|---|---|---|---|---|---|
| Crop | Image Format Conversion | Convert tiff image to mrc image | Eos | tiff2mrc | Pixel size of original image (Å) | ORGX=0.1019e4, ORGY=0.1019e4, ORGZ=0.209e4 | |
| | Resampling | Align x,y,z pixel spacing | Eos | mrcImageSamplingUnitChange | Pixel size after resampling (Å) | NEWX=0.1019, NEWY=0.1019, NEWZ=0.1019 | Interpolation: cubic interpolation method |
| | Checking the rotation angle | | IMOD or Fiji | | | | |
| | Image rotation | | Eos | mrcImageRotation3D | Rotation angle | ROTX=0.0, ROTY=31.5, ROTZ=0.0 | Apply padding with interpolation. |
| | | | | | Image size after resampling | ROTNX=840, ROTNY=512, ROTNZ=301 | |
| | Checking the crop position | | IMOD | | | | |
| | image cropping | | Eos | mrcImageCenterGet | Cropping position (center coordinates of the cropped image) | Set CENTERX, CENTERY, CENTERZ for each image. | |
| | | | | | Crop size | NX=256, NY=256, NZ=100 | |
| | Floating | | Eos | mrcImageFloating | | | |
| | Image Format Conversion | Convert mrc image to tiff image | Eos | mrc2tiff | | | raw image (32-bit) |
| Normalize | Image Format Conversion | | Eos | tiff2mrc | Pixel size of the cropped image | RESOLUTION=0.1019 | |
| | Normalization | Normalize by setting the High Value Area (0.001% of the total) to 0.7 and the Low Value Area (40% of the total) to 0.15. | Eos | mrcImageSeriesNormalizing | High Value Area | NORMAL_HIGH=0.99999 | |
| | | | | | Low Value Area | NORMAL_LOW=0.4 | |
| | | | | | High | NORMAL_A=0.70 | |
| | | | | | Low | NORMAL_B=0.15 | |
| | Image Format Conversion | Convert mrc image to tiff image | Eos | mrc2tiff | | | raw image (32-bit) |
| Creating mask images | Binarization | Set values below the background average of all frames (Low Value Area) + 6SD to 0, and values above that to 1. | Eos | mrcImageBinalization | threshold | The threshold value is calculated within the Makefile. | |
| | Calculation of cell area (volume) | Remove small noise in non-cellular areas. | Eos | mrcImageAreaCalc | | | |
| | Threshold setting | Set regions with a volume smaller than half of the cell's volume to 0. | Eos | mrcImageBinalization | threshold | The value obtained from MASKAREA is calculated within the makefile. | |
| | Apply a low-pass filter to the mask | | Eos | mrcImageLowPassFilter | Gauss filter | | Gaussian filter |
| | | | | | Gaussian function that decays to 1/e over 5 pixels | | |
| | Multiply the image by a value. | Multiply by 1/0.2 (or divide by 0.2) | Eos | mrcImageMultiplying | | The value to be applied to the image is calculated within the Makefile. | |
| | Binarization | Set values below the threshold to 0, and values equal to or above the threshold to 1. | Eos | mrcImageBinalization | threshold | 1 | |
| | Fill holes inside the mask. | 26-neighborhood connectivity | Eos | mrcImageHoleFilling | | | |
| | Smooth the edges. | Gradually transition the mask edges from 1 to 0. | Eos | mrcImageSoftEdge | width | MASKEDGE=5 | width: 5 pix |
| | | | | | envelop | MASKMODE=1 | Use a cosine function to transition from 1 to 0. |
| Creating a masked image | Add a value | Add a negative value to lower the overall values, ensuring the background becomes 0. | Eos | mrcImageAddValue | | | |
| | Masking | Apply the created mask to all images. | Eos | mrcImageMaskingByImage | | | -m 2 (accumulate mask image) |
| | Set the threshold | Set values below 0.07 to 0 (leave values above the threshold unchanged). | Eos | mrcImageBinalization | threshold | LASTTHRES=0.07 | |
| | | | | | value | LASTVAL=0 | |
| | Image format conversion | Convert to TIFF format. | Eos | mrc2tiff | | | raw image (32-bit) |
